# MyoD is a structure organizer of 3D genome architecture in muscle cells

**DOI:** 10.1101/2020.08.29.273375

**Authors:** Qian Chen, Fengling Chen, Ruiting Wang, Minglei Shi, Antony K. Chen, Zhao Ma, Guohong Li, Min Wang, Hu Li, Xu Zhang, Jinbiao Ma, Jiayun Zhong, Meihong Chen, Michael Q. Zhang, Yong Zhang, Yang Chen, Dahai Zhu

## Abstract

The genome is not a linear molecule of DNA randomly folded in the nucleus, but exists as an organized, three-dimensional (3D) dynamic architecture. Intriguingly, it is now clear that each cell type has a unique and characteristic 3D genome organization that functions in determining cell identity during development. A currently challenging basic question is how cell-type specific 3D genome structures are established during development. Herein, we analyzed 3D genome structures in primary myoblasts and myocytes from MyoD knockout (MKO) and wild type (WT) mice and discovered that MyoD, a pioneer transcription factor (TF), can function as a “genome organizer” that specifies the proper 3D genome architecture unique to muscle cell development. Importantly, we genetically demonstrate that H3K27ac is insufficient for establishing MyoD-induced chromatin loops in muscle cells. The establishment of MyoD’s “architectural role” should have profound impacts on advancing understanding of other pioneer transcription factors in orchestrating lineage specific 3D genome organization during development in a potentially very large number of cell types in diverse organisms.

## Introduction

The recent Hi-C analysis from comprehensive interaction map over large regions or whole genomes have indicated that the genome is hierarchically organized into chromosome territories, A/B compartments, topologically associated domains (TAD), and chromatin loops^1–3^. To date, much evidence has shown that the 3D structure of the genome is highly different among different cell types^4–6^, however, the molecular mechanisms underlying the establishment of cell-type specific 3D genome organization are largely unknown. It has been proposed that “pioneer transcription factors” act as anchor proteins to orchestrate cell-type specific 3D genome architecture^7,8^; however, there is no concrete genome-scale evidence to support this hypothesis. We selected MyoD for our exploration of pioneer transcription factors mediated specification of developmental-context specific 3D genome organization for the following reasons. First, MyoD is well-established as a pioneer transcription factor in myogenic cell lineage specification during development^9–12^ and trans-differentiation^13^. This protein is known to regulate the expression of myogenic specific genes through its binding to E-box (CANNTG) containing cis-regulatory elements^14,15^ and there are more than fourteen million consensus Eboxes in the genome^14^. Second, over forty thousand MyoD binding peaks identified by ChIP-seq from independent studies^15–17^ in muscle cells, only 15% of these MyoD binding peaks are located in promoter regions (Extended Data Fig. 1a). Finally, besides binding and transcriptional activation of genes during differentiation, MyoD also constitutively binds at tens of thousands of additional sites throughout the genome in proliferating muscle stem cells^15,18^ (Extended Data Fig. 1a). Together, these data represent an attractively large empirical scope to support a genome-wide functional analysis of MyoD’s potential roles in muscle cell lineage specific 3D genome organization. In this report, we provide computational and experimental evidence to uncover the unappreciated role of MyoD as a genome organizer in establishing the unique 3D genome architecture in muscle cells, beyond its well-known functions as a TF in activating myogenic gene expression during development.

## Results

### MyoD regulates A/B compartments switching and formation of contact domain boundaries (CDBs) in muscle cells

In light of the aforementioned MyoD data, we first performed RNA-seq analysis of proliferating (GM, Pax7^+^ and MyoD^+^) and differentiating (DM, MyoG^+^) muscle cells isolated from hind limb skeletal muscle of WT and MKO mice (Extended Data Fig. 1b), which we combined with MyoD ChIP-seq data to examine potential functional link(s) between MyoD occupancy and transcriptional activation. Specifically, we extracted MyoD binding peaks localized at promoter regions and identified their cognate genes within ± 3 kb (Fig. 1a): intriguingly, we found that knockout of MyoD did not affect the expression of a majority of these genes (76.7% for 2691/3510 in primary proliferation myoblasts, 56.6% for 1987/3509 in differentiation myocytes) (Fig. 1a and Supplementary Table 1), suggesting that the presence of MyoD at these genes (even in promoter regions) frequently does not directly impact their transcriptional activation. This result further supports the hypothesis that, beyond the canonical function of MyoD as an activator of myogenic gene expression during development, this protein may exert broad functional impacts, potentially on higher-order genome structures.

**Fig. 1.**
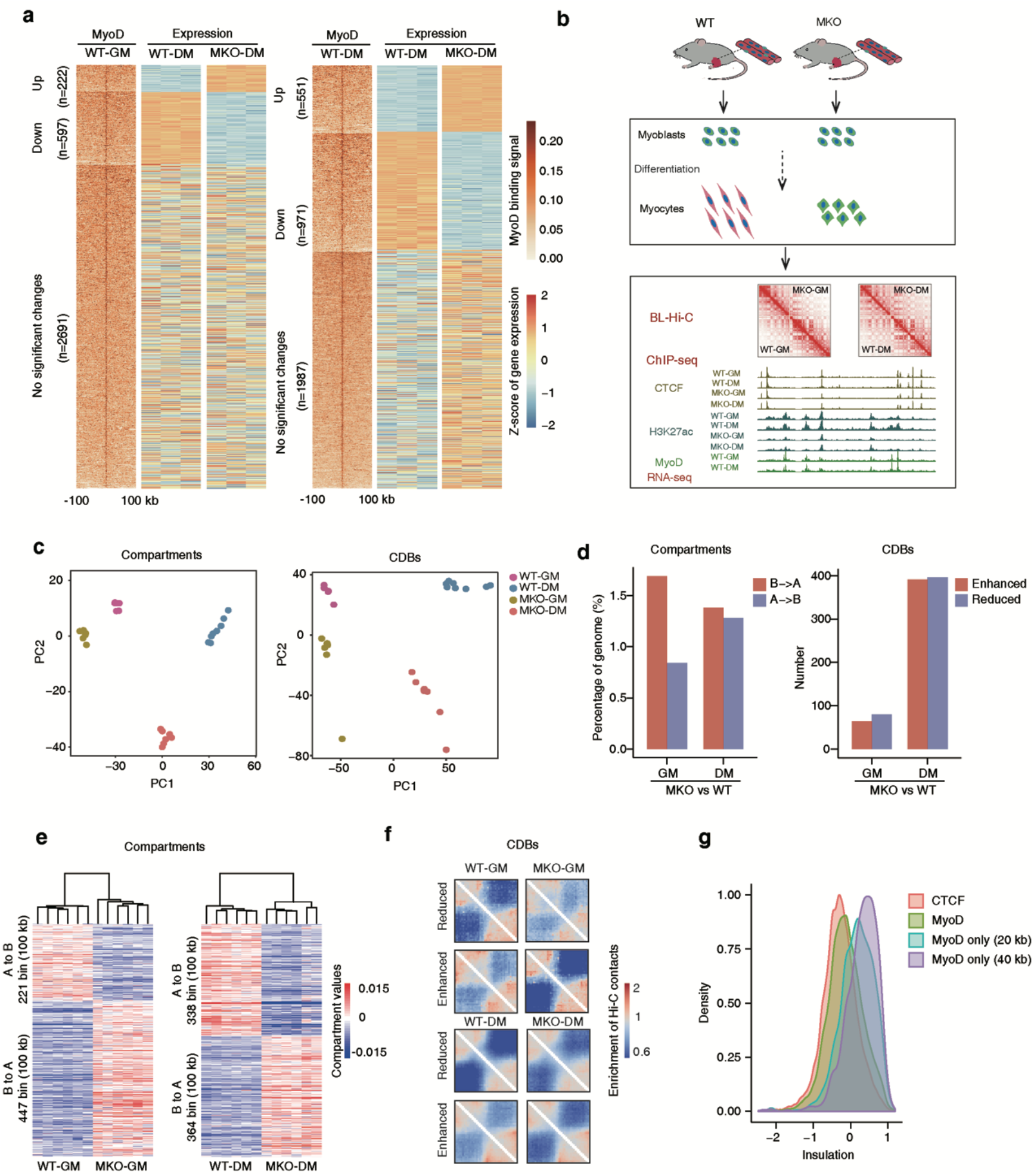
MyoD regulates A/B compartments switching and formation of contact domain boundaries (CDBs) in muscle cells. **a**, Association of MyoD binding peaks at promoter regions and related gene expression trends. Heatmaps showing MyoD binding peaks at promoters (centralized at the TSS within 3 kb distance) and related gene expression trends in WT and MKO myoblasts was shown at left panel. Corresponding data for WT and MKO myocytes was shown in the right panel. Pseudo-peaks of MyoD binding with combined ChIP-seq data from public databases and our study were used for this analysis (Methods). **b**, Strategy for genetic investigation of the chromatin architectural roles of MyoD. Briefly, muscle stem cells isolated from WT or MKO mice were cultured in GM and induced to differentiation in DM for 24 hrs. The proliferating myoblasts and differentiated myocytes were subjected to BL-Hi-C analysis. To correlate 3D genome architecture to chromatin state and to gene expression levels, multiple ChIP-seq and RNA-seq analyses were performed on myoblasts and myocytes from WT and MKO mice. **c**, Principal Component Analysis (PCA) of the values of compartments or the relative insulation scores of contact domain boundaries (CDBs) among the four indicated sample types. Each dot represents an individual biological replicate. **d**, Barplots showing the percentage of A/B compartment shifts between WT and MKO in myoblasts and myocytes (left panel). Barplots showing the number of relative insulation enhanced or reduced CDBs between WT and MKO in proliferating and differentiating muscle stem cells (right panel). **e**, Heatmaps of the A/B compartment shifted regions between WT and MKO in myoblasts and myocytes. **f**, Hi-C aggregation plots centered at differential CDBs between WT and MKO in myoblasts and myocytes. **g**, Probability density of insulation score at genetic regions bound by CTCF or MyoD. Four classes were assessed: CTCF-bound, MyoD-bound, MyoD-bound and CTCF-unbound within 20 kb distance, MyoD-bound and CTCF-unbound within 40 kb distance.

To directly investigate potential genome architectural roles of the pioneer TF MyoD in muscle cells, we further examined four muscle sample types (WT-GM, WT-DM, MKO-GM, MKO-DM) with bridge-linker Hi-C (BL-Hi-C)^19^ and ChIP-seq with antibodies against CTCF and histone mark H3K27ac (Fig. 1b). For the BL-Hi-C data, 28 libraries were sequenced to a total depth of over 12 billion reads; the high quality deep sequencing data were validated with high cis interaction rate (Extended Data Fig. 1c) and reached 5 kb resolution (Extended Data Fig. 1d), thereby ensuring the rigor of our subsequent computational analyses of each hierarchical level of chromatin structure in muscle cells.

We then examined the impacts of MyoD depletion on A/B compartments and contact domain boundaries (CDBs) in muscle cells by performing exploratory principal component analysis (PCA) based on the compartment values and relative insulation scores^20^ for the four muscle sample types examined (Methods). Our results showed clear distinctions between the WT and MKO genotypes for both proliferating and differentiating muscle cells (Fig. 1c). Moreover, we detected a clear A/B compartment shift in proliferating MKO cells vs. WT cells (1.69% B→A, 0.84% A→B) (Fig. 1d left and 1e left). For differentiating cells, there was a clear compartment switch for MKO cells compared to differentiating WT cells (1.38% B→A and A→B 1.28%) (Fig. 1d left, and 1e right).

We also analyzed CDB dynamics in muscle cells in response to MyoD knock out. A total of 931 CDBs exhibited differential extents of insulation in MKO cells compared to WT cells in proliferating and differentiating states (Fig. 1d right and 1f), implying that MyoD may somehow facilitate CDB insulation. Pursuing this, we compared insulation scores for CDBs bound by MyoD or by CCCTC-binding factor (CTCF). In general, the insulation scores of MyoD-bound CDBs were similar to the scores of CTCF-bound CDBs (Fig. 1g). However, the CDBs bound solely by MyoD exhibited significantly weaker insulation compared to the CTCF-bound CDBs, indicating that MyoD may work in concert with CTCF to regulate the insulation of CDBs in muscle cells.

Collectively, our analysis of the genome-wide distribution of MyoD alongside functional studies of MKO proliferating and differentiating muscle cells support the idea that MyoD has previously unappreciated functional roles in regulating A/B compartments and contact domain boundaries in muscle cells.

### MyoD is an anchor protein for chromatin loop formation in muscle cells

The most well-characterized anchor protein in vertebrates is CTCF, which mediates chromatin looping in concert with cohesin probably through the loop extrusion mechanism^2,21–25^. Considering our finding that MyoD apparently functions together with CTCF to regulate insulation of CDBs, we wondered whether MyoD may control the formation of chromatin loops in muscle cells. Using HiCCUPS algorithm^26^ to assess our BL-Hi-C dataset, we ultimately identified 24,492 chromatin loops in proliferating WT myoblasts and 31,243 chromatin loops in differentiating WT myocytes (Fig. 2a and Supplementary Table 2, Methods). The accuracy of the loop calls was supported by high-scoring aggregate peak analysis (APA) plots (Fig. 2a, Methods) and by observable enrichment of CTCF-bound loops in muscle cells (Extended Data Fig. 2a) wherein about 75% of them had convergent CTCF binding motifs (Extended Data Fig. 2b).

**Fig. 2.**
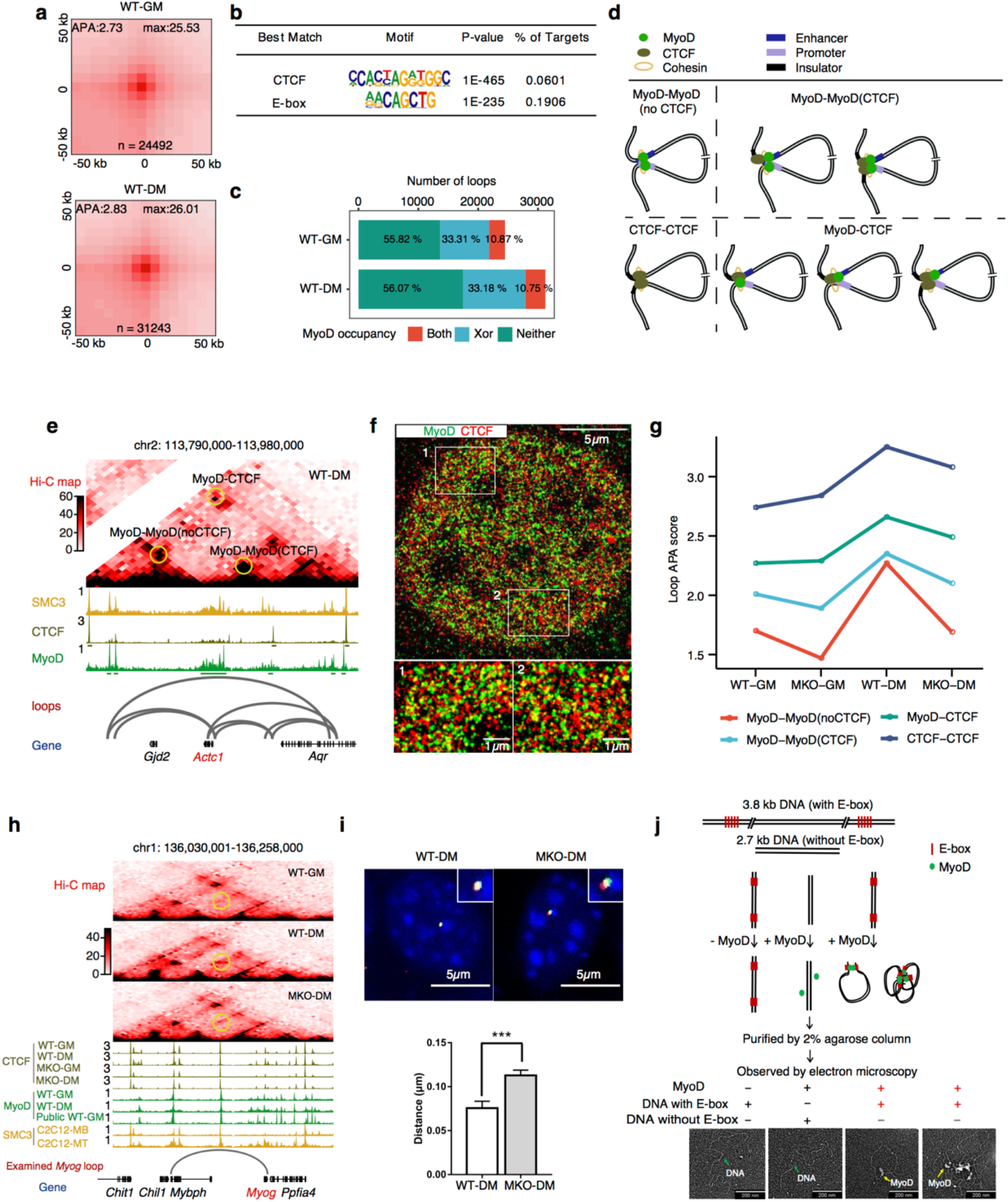
MyoD is an anchor protein for chromatin loop formation in muscle cells. **a**, Aggregate Peak Analysis (APA) plots showing the aggregated Hi-C contacts around chromatin loops identified in WT-GM and WT-DM cells. n represents the total number of chromatin loops called by HiCCUPS (Methods). APA score at upper left corner of each aggregation plot was calculated as the ratio of Hi-C contacts at the central pixel to the mean Hi-C contacts of the lower left pixels. The Hi-C contacts at the central pixel was shown at upper right corner of each aggregation plot. **b**, Enriched motifs in accessible regions at anchors of the chromatin loops identified in panel a. **c**, Percentage of chromatin loops with anchors bound by MyoD in WT-GM and WT-DM cells. MyoD bound at both anchors (Both, red), one of two anchors (Xor, blue), or neither of two anchors (Neither, green). Pseudo-peaks of MyoD binding with combined ChIP-seq data from public databases and our study were used for this analysis (Methods). **d**, Classification of the four types of chromatin loops identified in the study. MyoD-MyoD(noCTCF) loops had both anchors bound only by MyoD. MyoD-MyoD(CTCF) loops had both anchors bound by MyoD and either anchor concomitantly bound by CTCF. MyoD-CTCF loops had only one anchor bound by MyoD and either anchor bound with CTCF. CTCF-CTCF loops had both anchors bound only by CTCF. **e**, Representative examples of the MyoD-bound chromatin loops. ChIP-seq of SMC3 were in C2C12 cells referenced from public database (GSE113248). **f**, Representative dSTORM image showing co-localization of MyoD (green) and CTCF (red) from immunofluorescent staining of C2C12 cells grown in DM for 24 hrs with antibodies against MyoD and CTCF. **g**, APA score dynamics for the four types of chromatin loops among WT and MKO cells in GM and DM stages. **h**, A representative MyoD-bound chromatin loop at the *Myog* locus that presents the Hi-C contact map depicting normalized contact frequencies and ChIP-seq signal tracks. The examined chromatin loop with reduced Hi-C contacts in MKO-DM sample (compared with WT-DM sample) was marked by yellow circle. **i**, Chromatin loop at the *Myog* locus was validated by 3D-DNA fluorescent *in situ* hybridization (3D-DNA FISH) in WT-DM and MKO-DM cells. The upper panel showed *Myog* promoter (red) and enhancer (green) were more separated in the MKO cells (right) than in the WT cells (left). The lower panel showed the average distances of *Myog* loop anchors. ***P < 0.001, two-tail student’s t-test. **j**, MyoD mediates chromatin looping *in vitro*. The upper panel presents a schematic for the *in vitro* DNA circularization assay used to detect the ability of MyoD (green) to mediate DNA looping with E-box (red) containing DNA (3.8 kb) and no E-box control (2.7 kb). The lower panel presents representative views of circularized DNA mediated by recombinant MyoD protein, imaged by Transmission Electron Microscopy (TEM).

We then searched for enriched motifs within accessible regions in loop anchors and detected significant enrichment for known myogenic E-box motifs^14,15^ (Fig. 2b). This enrichment was further confirmed by our observation of enrichment for MyoD binding peaks at loop anchors as analyzed using pseudo-peaks (Methods) with combined MyoD ChIP-seq data from our present study and other published work^17^ (Fig. 2c and Supplementary Table 3): 44.18% of chromatin loops in GM cells were MyoD-bound (10.87% with both anchors bound by MyoD, 33.31% with one anchor bound by MyoD), while 43.93% of chromatin loops in DM cells were MyoD-bound (10.75% with both anchors bound by MyoD, 33.18% with one anchor bound by MyoD). Furthermore, MyoD binding peaks at loop anchors were concordant with binding signals of CTCF and SMC3^27^, a subunit of the cohesin complex (Extended Data Fig. 2c). We cataloged all of the identified chromatin loops into four types based on the binding of MyoD and/or CTCF at loop anchors in muscle cells: MyoD-MyoD(noCTCF), MyoD-MyoD(CTCF), MyoD-CTCF and CTCF-CTCF (Fig. 2d, e). This classification was further supported by the observable MyoD co-localization with CTCF as well as non-overlaid MyoD signals or CTCF signals when the two proteins were imaged by direct stochastic optical reconstruction microscopy (dSTORM), a single-molecule localization microscopy technique that can reveal the organization of specific proteins with a lateral resolution of ~20 nm^28^ (Fig. 2f). In addition, we found that the MyoD-bound loops were significantly shorter than CTCF-CTCF loops (288 kb on average), and the MyoD-MyoD(noCTCF) chromatin loops were even shorter (113 kb on average) (Extended Data Fig. 2d). Together, these data substantiate MyoD localization on anchors of chromatin loops in muscle cells.

We next assessed whether MyoD could indeed mediate chromatin loop formation *in vivo* by investigating the effects of MyoD knockout on the MyoD-bound loops detected in WT cells. APA plots indicated that MyoD knockout significantly decreased the loop strength values for both MyoD-bound chromatin loops (MyoD-MyoD(noCTCF), MyoD-MyoD(CTCF) and MyoD-CTCF loops) and CTCF-CTCF chromatin loops, with these decreases evident in both GM and DM cells (Fig. 2g and Extended Data Fig. 2e). These results support that MyoD functionally contributed to the formation of both MyoD-bound and CTCF-bound chromatin loops *in vivo*. Consistent with this possibility, fluorescence *in situ* hybridization (FISH) confirmed that the MyoD-bound chromatin loop connecting the *Myog* promoter and the *Mybph* promoter was established at the *Myog* locus in differentiating WT cells but not in differentiating MKO muscle cells (Fig. 2h, i). Moreover, *in vitro* DNA circularization assays with DNA fragments bearing known MyoD binding motifs (Methods) showed that MyoD protein induced the circularization of linear DNA with MyoD binding motifs *in vitro* (Fig. 2j and Extended Data Fig. 2f, g), confirming MyoD-orchestrated chromatin looping.

Collectively, these genetic, biochemical, and cellular imaging results demonstrate that MyoD functions as an anchor protein that orchestrates chromatin loop formation across the genome in muscle cells. Moreover, our data support that MyoD impacts chromatin loop formation by itself and through its physical interaction with CTCF.

### MyoD anchors formation of myogenic-specific chromatin loops

Based on the above findings, we next asked whether MyoD can mediate the organization of muscle cell-specific chromatin looping. To test this, Hi-C data from embryonic stem (ES) and cortical neuron (CN) cells^5^ were collected, and the same four types of chromatin loops identified in muscle cells were examined in ES and CN cells. Only a few of MyoD-MyoD(noCTCF) loops were identified in ES cells (only 8 loops, 1.8%) or CN cells (only 1 loop, 0.2%) (Fig. 3a), revealing that the MyoD-MyoD(noCTCF) loops detected in muscle cells represent myogenic lineage specific chromatin loops. Strikingly, by assessing gene expression trends associated with four types of loops across eight cell types (Methods), we demonstrated that the genes from the MyoD-MyoD(noCTCF) loops exhibited the strongest extent of muscle lineage specific expression (Fig. 3b). Accordantly, the genes associated with MyoD-MyoD(noCTCF) chromatin loops also had the most enriched GO terms related to muscle cell differentiation (Extended Data Fig. 3a).

**Fig. 3.**
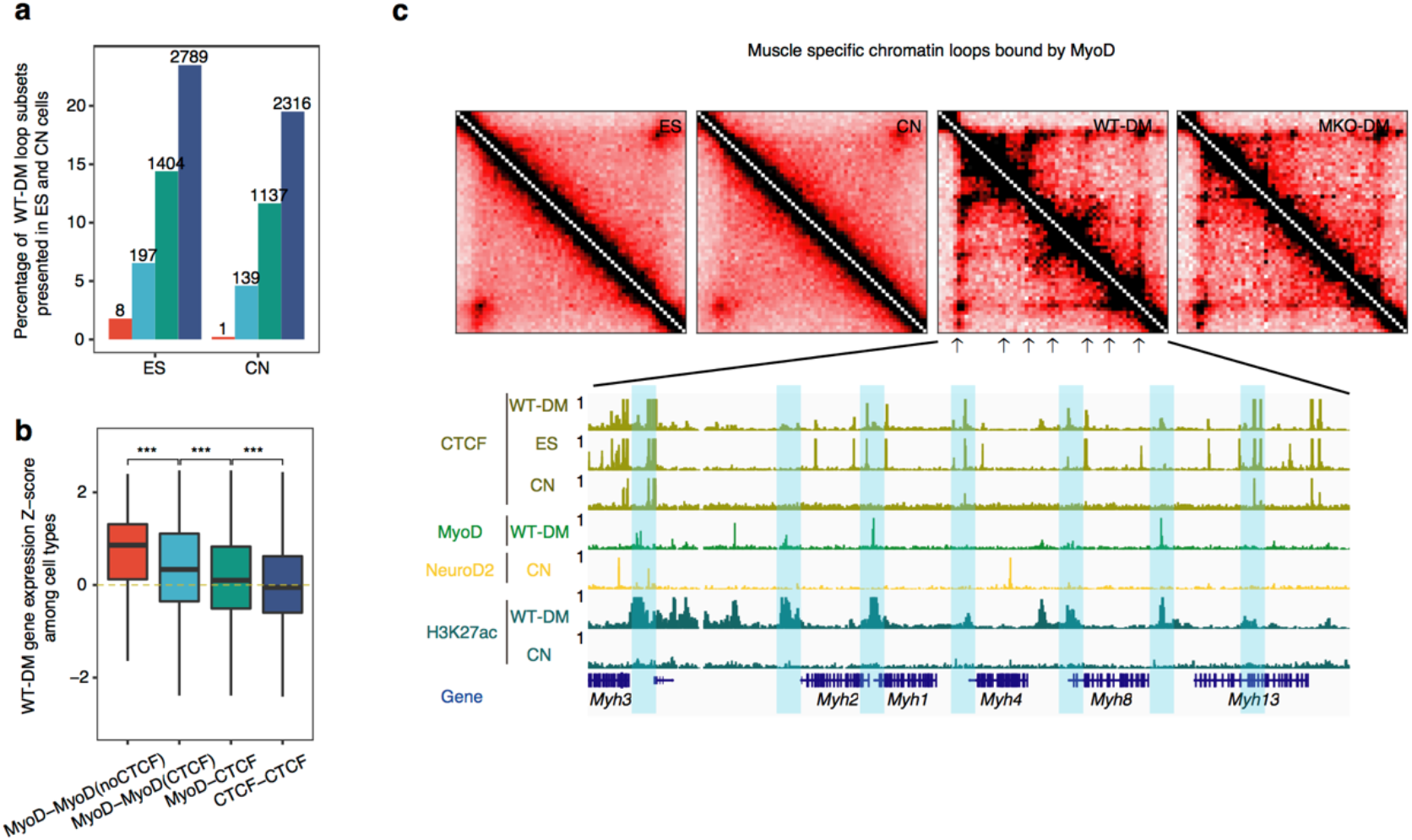
MyoD anchors formation of myogenic-specific chromatin loops. **a**, Proportions of the muscle chromatin loop subsets presented in embryonic stem cell (ES) or cortical neuron (CN). The numbers of muscle chromatin loops found in ES or CN cells were indicated for each bar. **b**, Boxplots represent Z-score of gene expression in WT-DM cells compared with other ENCODE collected cell types (embryonic stem cell, spleen, B cell, T cell, megakaryocyte, neural progenitor and cortical neuron) for genes associated with each class of loops, showing MyoD-bound loops regulate lineage specific gene expression in muscle cells. ***P < 0.001, Wilcoxon rank sum test. **c**, Representative region showing muscle cell specific loops anchored on MyoD binding regions at *Myh* gene cluster.

To seek further molecular insights into MyoD-bound muscle cell-specific chromatin looping, we examined ChIP-seq data for both MyoD and NeuroD2 — a pioneer TF for neuronal cell lineage commitment and differentiation^18,29^ in both muscle cells and neuron cells. Only 18% of MyoD binding peaks in muscle cells were overlapped with NeuroD2 binding peaks in cortical neuron cells, suggesting that MyoD and NeuroD2 might mediate the formation of separate sets of chromatin loops that can subsequently regulate distinct sets of target genes to properly specify lineage-specific cell identities. For example, we could identify that the *Myh* gene cluster that are located within MyoD-bound, muscle-cell-specific chromatin loop were not detected in neuronal cells, and there was no NeuroD2 binding at the same genomic region in neuronal cells (Fig. 3c). *Vice versa*, chromatin loops bound by the NeuroD2 at the *Foxg1* loci is neuron-specific and not observed in muscle cells (Extended Data Fig. 3b). Taken together, these results illustrate that cell-lineage pioneer TFs exert genome-wide architectural roles for modulating 3D genome organization, a functional impact considerably larger than the previous, primarily gene-transcriptional-activation related understanding about the influence of pioneer TFs on lineage specification.

### MyoD mediates primed architectural chromatin loops in proliferating muscle cells

Recalling previously published studies showing that MyoD constitutively binds to tens of thousands of sites genome-wide in proliferating cells^15,18^, together with our present finding that such binding only infrequently results in transcriptional activation at this stage, and that MyoD functions as an anchor protein for lineage-specific 3D genome organization, we sought to further explore the nature of MyoD’s genome architectural functions by re-assessing our BL-Hi-C data. We were particularly interested in addressing the question of how the presence of MyoD influences internal interactions occurring within chromatin loops.

Previous studies have indicated that CTCF and cohesin function together to establish architectural loops and that the extent of enhancer-promoter (E-P) interactions is elevated within such loops^30^. As our results identify a notable co-localization of MyoD binding peaks with CTCF (Fig. 2f), we speculated that MyoD-bound loops may also feature increased intraloop interactions. To test this, we measured internal interactions for each loop by calculating “domain score” (D-score) in WT and MKO proliferating cells. MyoD binding is enriched on anchors of chromatin loops with reduced internal interactions (65%) in MKO-GM cells compared with WT-GM cells (Fig. 4a, b), indicating that MyoD regulates the internal interactions within chromatin loops by its direct binding on loop anchors like CTCF and cohesin. Further analysis showed that interactions changes within loops were positively correlated with expression changes of genes enclosed in the same chromatin loop (Fig. 4c).

**Fig. 4.**
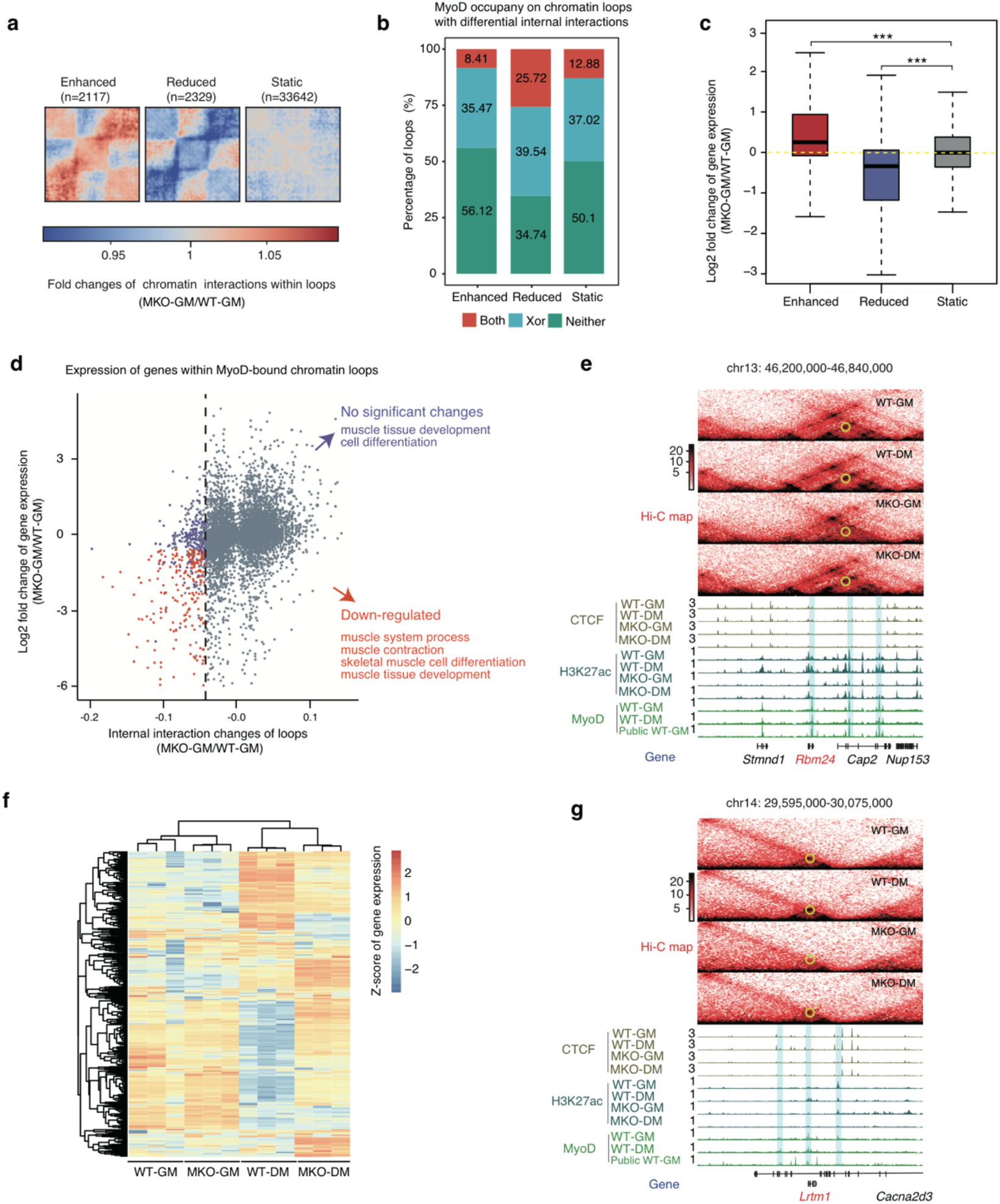
MyoD mediates primed architectural chromatin loops in proliferating muscle cells. **a**, Aggregated Hi-C maps around chromatin loops with enhanced, reduced, or static internal interactions (measure by domain score) between WT and MKO myoblasts (MKO-GM vs. WT-GM). **b**, Percentage of chromatin loops with differential internal interactions (MKO-GM vs. WT-GM) that occupied by MyoD at anchors. MyoD bound at both anchors (Both, red), one of two anchors (Xor, blue), or neither of two anchors (Neither, green). Pseudo-peaks of MyoD binding with combined ChIP-seq data from public datasets and our study were used for this analysis (Methods). **c**, Boxplot showing the corresponding expression changes of genes within chromatin loops having enhanced, reduced, or static internal interactions (domain score) identified in panel b (MKO-GM vs. WT-GM). Considering that one loop may contain several genes and domain scores of nested loops are associated, we assigned each gene to its most dynamic nested loop. ***P < 0.001, Wilcoxon rank sum test. **d**, Scatter plot showing both the domain score fold changes and gene expression fold changes of MyoD-bound chromatin loops between WT-GM and MKO-GM cells. Internal interaction reduced MyoD-bound chromatin loops without significantly differentially expressed genes in MKO-GM cells compared with WT-GM cells were shown as blue dots, while internal interaction reduced MyoD-bound chromatin loops with significantly down-regulated genes in MKO-GM cells compared with WT-GM cells were shown as red dots. **e**, The *Rbm24* locus as a representative for an internal interaction reduced MyoD-bound chromatin loop with significantly down-regulated genes in MKO-GM cells compared with WT-GM cells. **f**, Heatmap showing expression (FPKM) Z-scores among four indicated sample types of genes within internal interaction reduced MyoD-bound chromatin that do not contain differentially expressed genes in MKO-GM cells compared with WT-GM cells. **g**, The *Lmrt1* locus as a representative for an internal interaction reduced MyoD-bound chromatin loop in MKO-GM cells compared with WT-GM cells that is primed in proliferating muscle cells.

We next focused on analyzing the internal interaction changes of MyoD-bound loops and their cognate gene expression (Fig. 4d). We specifically examined those MyoD-bound loops displaying reduced interactions in MKO proliferating cells because these loops might be regulated directly by MyoD binding. We found that nearly 50% of the examined loops enclosed down-regulated genes with known myogenic-related functions (Fig. 4d), indicating that interactions within these MyoD-bound loops are necessary for the regulation of muscle identity gene expression. For example, the *Rbm24* locus, a known gene that regulates myogenesis^31^ and its differential expression in WT-GM and MKO-GM cells offers an excellent illustration for a gene enclosed within this type of MyoD-bound loop in muscle cells (Fig. 4e and Extended Data Fig. 4a).

Intriguingly, we also found that most (89%) of the remaining MyoD-bound loops with decreased interactions did not contain any differentially expressed genes upon MyoD knockout (Fig. 4d). This finding underscored the apparent centrality of the architectural role (rather than transcriptional activation) of MyoD for these MyoD-bound loops in proliferating muscle cells. Although reduced interactions within these chromatin loops in MKO cells did not alter gene expression in proliferating cells, we found that genes within these chromatin loops were not turned on or off properly in the MKO-DM cells as compared with WT-DM cells (Fig 4f), and this genetic dysregulation was clearly manifest in the defective differentiation phenotype of the MKO cells (Extended Data Fig. 1b) ^32–34^ As the loops were still maintained in differentiated cells (Extended Data Fig. 4b), it appears that the interactions constrained within the loops in proliferating cells are effectively “primed” for muscle cell differentiation by the presence of MyoD, raising the possibility that some very early signal can somehow direct the occupancy of MyoD to the sites across the genome that well subsequently permit both muscle-cell-appropriate loop architecture and later rewiring in response to differentiation signals. Supporting this idea, we found that *Lrmt1*, a known myogenesis related gene^35^, which is only expressed in differentiated muscle cells (Extended Data Fig. 4c), contains chromatin loops that have already been defined by MyoD in proliferating cells (Fig. 4g). Thus mechanistically, our analysis of the MyoD-bound loops and their internal interactions revealed molecular insights about how MyoD exerts its architectural function, even in proliferating cells, to ensure the correct trajectory towards eventual muscle cell identity.

### The regulatory loops specified by MyoD are functionally required for muscle cell differentiation

The genome is not only structurally organized within the nucleus but is also dynamically orchestrated in response to various cellular signals^5,36–40^. However, the molecular mechanisms underlying dynamic changes in 3D genome structure are largely unknown. Although the biological function of MyoD in regulating muscle cell differentiation has been well documented, and very recently, MyoD’s transcription activity was found to be regulated by its ability to mediate chromatin interactions during trans-differentiation^41^, how MyoD impacts chromatin conformation dynamics during muscle cell differentiation remains largely elusive. Using a modified differential loop detection method^40^ (Methods), we identified 6,242 differential chromatin loops (~25%) between proliferating and differentiating muscle cells. Among these loops, 5,754 were significantly enhanced whereas 488 were reduced in differentiating cells (Fig. 5a), and we detected significant enrichment for muscle cell differentiation related genes associated with the enhanced chromatin loops (Extended Data Fig. 5a). These data also revealed that nearly 25% of chromatin loops that undergo dynamic changes during muscle cell differentiation contribute to 3D genome rewiring in a manner that functionally regulates myogenic differentiation.

**Fig. 5.**
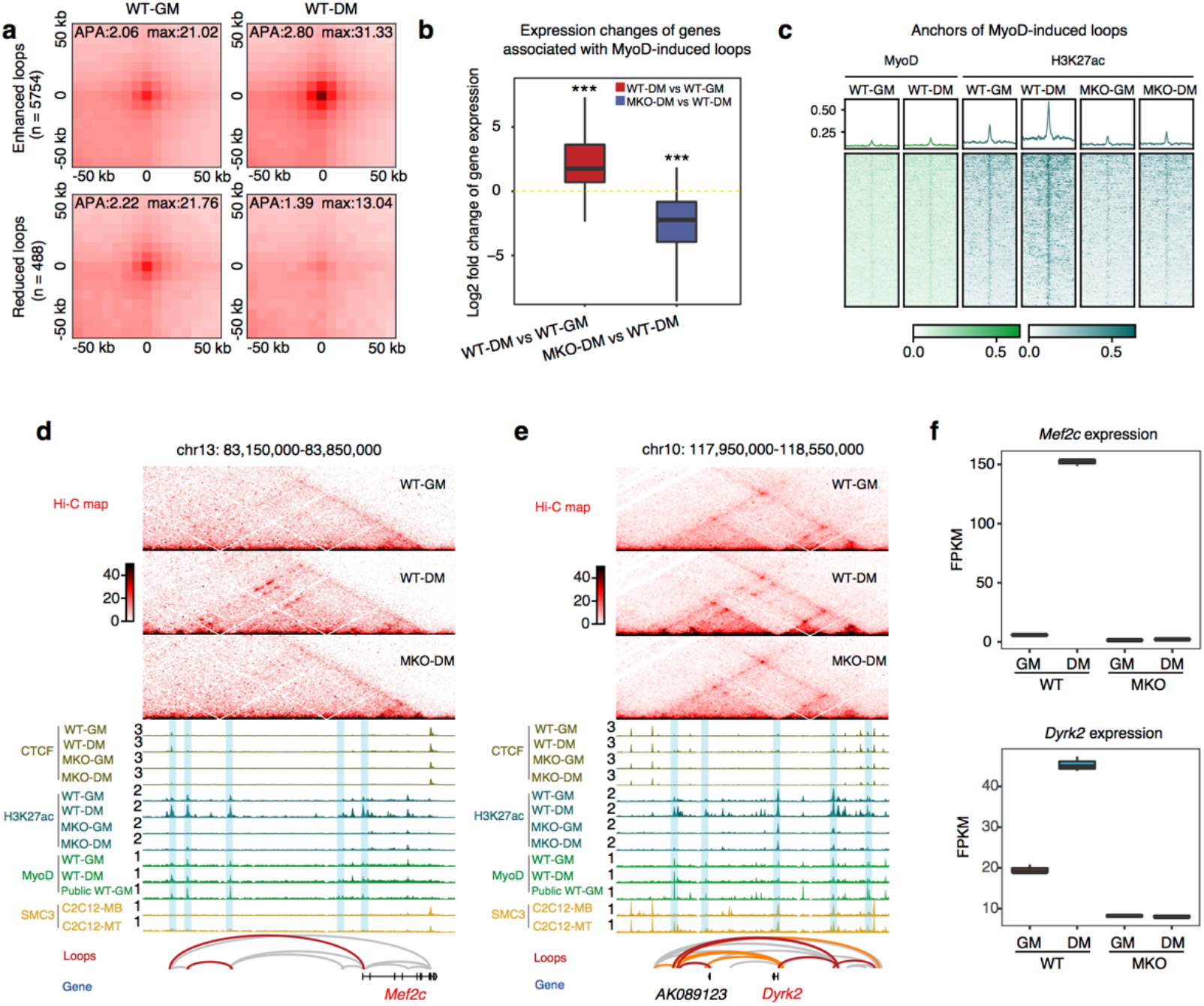
The regulatory loops specified by MyoD are functionally required for muscle cell differentiation. **a**, APA plots showing the aggregated Hi-C contacts around enhanced (n=5754) and reduced (n=488) chromatin loops in WT-DM cells compared with WT-GM cells. Differential loops were identified based on DESeq2 (Methods). **b**, Boxplot showing expression changes of genes associated with MyoD-induced chromatin loops in WT-DM cells compared with WT-GM cells (left) and in MKO-DM cells compared with WT-DM cells (right). ***P < 0.001, Wilcoxon rank sum test. **c**, MyoD and H3K27ac ChIP-seq signal at anchors of MyoD-induced loops. Heatmaps showing signal enrichment centralized at the anchors within ± 100 kb genomic region. **d** and **e**, The*Mef2c* and *Dyrk2* locus as representatives of MyoD-induced loops during muscle cell differentiation. MyoD-induced loops detected by our algorithm were presented in red in the loop track while differentiation-induced loops were in orange. Other loops were presented in grey. **f**, *Mef2c* and *Dyrk2* gene expression among WT and MKO cells in proliferating and differentiating stage.

Moreover, similar analysis performed on MKO cells identified 585 differential chromatin loops between WT-DM and MKO-DM cells. Consistent with their regulatory role, 81.5 % (477/585) of these were significantly reduced in the differentiated MKO cells (Extended Data Fig. 5b), and 71.57% (341/477) of the reduced loops identified in the differentiated MKO cells had MyoD binding peaks support them as bona fide MyoD mediated chromatin loops in differentiated WT cells (Extended Data Fig. 5c). Among the aforementioned enhanced loops in differentiation, 285 loops might be directly induced by MyoD as they were bound by MyoD and disappeared when MyoD was absent in differentiating muscle cells (Supplementary Table 4). The genes associated with those MyoD-induced loops significantly up-regulated during differentiation and down-regulated when MyoD was absent, indicating the regulatory role of MyoD-induced loops (Fig. 5b).

Next, we sought to understand the driving force for the enhancement of MyoD-induced loops during muscle differentiation. One possibility might be due to increased MyoD binding on loop anchors during muscle cell differentiation. By examining MyoD binding dynamics on anchors of MyoD-induced loops, we found that majority of MyoD-induced loops did not show novel MyoD binding during the differentiation (Fig. 5c). However, we found that 88% of MyoD-induced loops were decorated with MyoD dependent H3K27ac modification (Fig. 5c). Collectively, these findings raise possibility that H3K27ac may contribute to loop formation during muscle cell differentiation and that MyoD somehow actively directs the deposition of these modifications on the MyoD-induced loops.

Taking the *Mef2c* locus as an example of MyoD-induced loops to demonstrate their regulatory role in differentiation, we found that enhancer-promoter loops linked to *Mef2c* emerged during differentiation in a MyoD dependent manner (Fig. 5d). Concordantly, H3K27ac modification was enhanced in WT-DM cells but diminished in MKO-DM cells at this locus (Fig. 5d) while *Mef2c* gene expression was not induced in MKO-DM cells (Fig. 5f upper). Similar results were obtained when we examined MyoD-induced loops within the *Dyrk2* locus, from which the mRNA transcription product was only abundant in WT-DM cells (Fig. 5e and Fig. 5f lower). Collectively, these results demonstrate MyoD orchestrates the formation of myogenic lineage specific chromatin loops that are required for the transcriptional regulation of myogenic genes during muscle cell differentiation.

### H3K27ac in the MyoD knock out cells is not sufficient to drive formation of the MyoD-bound chromatin loops

The dynamic organization of the 3D genome is known to facilitate sophisticated interplay between aaccessible versus inaccessible chromatin states and TF occupancy^38^; however, the interdependent impacts (and likely feedback) of chromatin status on 3D genome organization have remained largely elusive. Aside from MyoD dependent H3K27ac modification on MyoD-induced loops, we found that during muscle cell differentiation, the loop anchors which had increased H3K27ac signals showed enrichment for consensus E-box sequences (i.e., the known target motif of the MyoD protein) in general (Fig. 6a). When MyoD was knocked out, H3K27ac modification was decreased on MyoD binding regions (Fig. 6b) which further support the overall dependency of H3K27ac modification on MyoD during differentiation. As MyoD itself stably binds around many myogenic genes in proliferating muscle cells, MyoD dependent H3K27ac modification correlates even better with the loop formation or disruption. These results and previously reported positive correlation between H3K27ac and loop^39,42^ prompted us to examine whether H3K27ac-mediated active chromatin state directly triggers the formation of MyoD-bound chromatin loops during muscle cell differentiation.

**Fig. 6.**
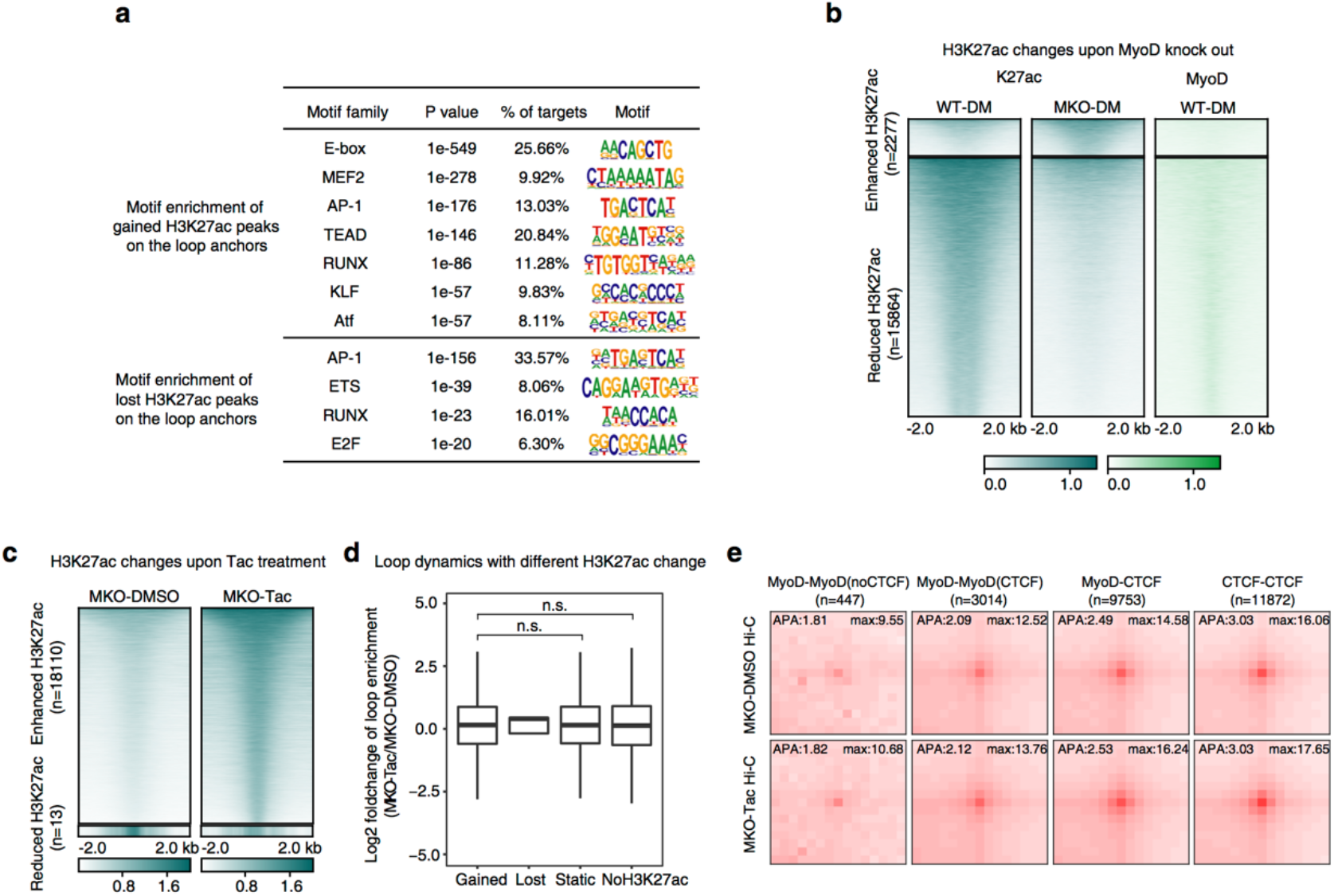
H3K27ac in the MKO cells is not sufficient to drive formation of the MyoD-bound chromatin loops. **a**, Motif enrichment at gained or lost H3K27ac peaks on the loop anchors in WT-DM cells compared with WT-GM cells. **b**, Heatmaps showing enhanced and reduced H3K27ac peaks from ChIP-seq analysis in MKO-DM cells compared with WT-DM cells. MyoD binding peaks at those differential H3K27ac peaks were also shown. **c**, Heatmaps showing enhanced and reduced H3K27ac peaks from ChIP-seq analysis in MKO cells treated with a selective class I HDAC inhibitor Tacedinaline (Tac) in differentiation medium for 12 hrs. DMSO treated MKO cells served as control. **d**, Loop strength fold changes of the loops with gained, lost, static or no H3K27ac marks on their anchors between MKO cells treated with Tac or DMSO. Loop strength was defined as relative Hi-C contacts at loop pixels compared with surrounding Donut-shaped background pixels (Methods). ***P < 0.001, Wilcoxon rank sum test. **e**, APA plots showing aggregated Hi-C contacts around the four types of chromatin loops (classified in 2d) in MKO cells treated with Tac and DMSO control.

We tested this possibility by treating MKO cells with the histone deacetylase I specific inhibitor Tacedinaline (Tac) to enrich the H3K27ac signal in the cells throughout muscle cell differentiation. It was found that Tac treatment could result in the accumulation of H3K27ac modification in 18, 110 genomic regions including anchors of MyoD-induced loops (96/285) (Fig. 6c). Furthermore, analysis of the BL-Hi-C data from the Tac-treated MKO and DMSO-treated cells showed that, in the absence of MyoD, the accumulation of H3K27ac *per se* is not sufficient to trigger the formation of new loops or to enhance the signal for MyoD-bound or CTCF-CTCF chromatin loops (Fig. 6d, e). Thus, H3K27ac alone is insufficient for establishing characteristic 3D genome organization in muscle cells, as simply accumulating H3K27ac cannot rescue MyoD-induced loops or drive other loop formation. Together, our overall findings reveal a critically determination role of the pioneer TF MyoD as a genome organizer in establishing the unique 3D genome architecture in muscle cells.

## Discussion

Each cell type acquired unique stage-specific organization of 3D genome during development and cell-type specific organization of 3D genome can be regarded as an emergent property that is mediated by interplay between transcription factors and chromatin-associated proteins^43^. Based on this model, the components in regulating the emergent property of chromatin should be coordinately orchestrated by a genome organizer to instruct specific organization of 3D genome in each cell type at a specific developmental stage. In this report, we provide compelling evidence to support this model wherein binding of the pioneer transcription factor MyoD at sites across the genome in muscle stem cells exerts previously unappreciated function as a genome organizer, which has broad functional impacts on genome structure beyond its known functions in activating gene expression during development (Extended Data Fig. 6). Mechanistically, we show that internal interaction changes in MyoD-bound loops suggest a new insight into the potential mechanism of cell lineage pioneer TFs to ultimately specify the positions and nature of the chromatin loops comprising the cell-type specific internal interactions which ensures the subsequent differentiation process leading to lineage specific cell identity. Further, we biochemically demonstrate that interaction between a pioneer TF MyoD and structural protein CTCF drives different types of loops which ultimately supports a lineage specific genetic programme. Finally, we show how the occupancy of pioneer TF proteins coordinate the epigenetic mark H3K27ac to retain active expression of the pioneer-TF-specified genome structures and developmental genetic programmes underlying proper muscle stem development.

It will be interesting to determine which signals ultimately initiate MyoD’s loop specification layer of genetic regulation at the earliest possible stage of myogenic lineagespecific cell fate determination. Excitingly, extending our insights beyond MyoD-mediated architectural regulation of muscle cell fate, we have also provided additional evidence from experiments with the well-studied neuron specific pioneer TF NeuroD2 in neuronal cells. This result suggests a highly similar loop specification function of NeuroD2 which apparently underlies the neuron lineage specific 3D genome architecture and attends (downstream) genetic programming of future cell fate. We are eager to see if other lineage-specific pioneer TFs in different developmental systems or organisms could confirm the general role, similar to that played by MyoD in our discovery, as the genome organizers that orchestrate the lineage specific chromatin and 3D genome structures required for cell fate determination during development.

## Acknowledgments

This work was supported by grants from the National Key Research and Development Program of China (2016YFA0100703, 2017YFA0505503), the National Natural Science Foundation of China (Nos. 91540206, 31871343, 31671384, 31971080 and 91949106), the Natural Science Foundation in Beijing (No. 7192125), and the CAMS Initiative for Innovative Medicine (Nos. 2016-I2M-1-017, 2019-I2M-1-004). The funding agencies do not involve in any research activities in the present study.

We thank Professor Cheng Li and Professor Bing Zhu for useful discussion and critical reading of the manuscript.

## Author contributions

Q.C. conducted cell culture, BL-Hi-C, ChIP-seq, RNA-seq libraries and wrote the manuscript. F.C. performed all computational and bioinformatics analysis, generated the figures and tables as well as wrote the manuscript. R.W. carried out 3D-FISH analysis and immunostaining of MyoD and CTCF. M.S. helped to perform BL-Hi-C libraries. M.C. helped data analysis. A.K.C. and Z.M. performed dSTORM analysis for co-localization of MyoD and CTCF. G.L. and M.W. conducted in vitro DNA circularization and transmission electron microscopy. H.L. helped data analysis. X.Z. helped to conduct 3D-FISH. J.M. and J.Z. purified recombinant MyoD protein. Y.C. and M.Q.Z. supervised computational and bioinformatics analysis. Y.Z. helped to design and supervised the experiments and wrote the manuscript. D.Z. conceived of and supervised the project and wrote the manuscript. The authors declare no competing financial interests.

## Methods

### Cell line

#### C2C12 cell culture and differentiation

Mouse C2C12 cells (ATCC, CRL-1772) were cultured in growth medium consisting of Dulbecco’s modified Eagle’s medium (Gibco) supplemented with 4.5 g/L glucose, 10% fetal bovine serum (Ausbian), 1% penicillin and streptomycin at 37°C in a 5% CO_2_ atmosphere. For the differentiation of C2C12 myoblasts, cells were transferred to Dulbecco’s modified Eagle’s medium containing 2% horse serum and 1% antibiotics, and then cultured for 24 hrs. C2C12 cells were grown to 80–90% confluence before induction of differentiation.

### Mouse strains

*MyoD KO* mice (#002523) were obtained from the Jackson Laboratory. All the animal procedures were approved by the Animal Ethics Committee of Peking Union Medical College, Beijing, China (ACUC-A01-2016-003). Mice were housed in the animal facility and had free access to water and standard rodent chow.

### Primary myoblasts

#### Primary myoblast isolation, culture and differentiation

Primary myoblasts were isolated from hind limb skeletal muscle of MKO and WT littermates at 2~3 weeks old, minced, and digested in a mixture of type II collagenase and Dispase. Cells were filtered from debris, centrifuged, and cultured in growth medium (F-10 Ham’s medium supplemented with 20% fetal bovine serum, 10 ng/ml basic fibroblast growth factor and 1% antibiotics) on collagen-coated cell culture plates at 37°C in 5% CO_2_. For the differentiation of primary myoblasts, cells were transferred to Dulbecco’s modified Eagle’s medium (Gibco) containing 2% horse serum and 1% penicillin and streptomycin, and then cultured for 24 hrs. All cells were grown to 60–70% confluence before induction of differentiation. For H3K27ac accumulation experiments, MKO myoblasts were treated with 2.5 μM Tacedinaline (CI994) in differentiation medium for 12 hrs to enrich H3K27ac while DMSO treated MKO myoblasts in DM for 12 hrs served as control. Then the cells were collected for ChIP-seq and BL-Hi-C. Tacedinaline (CI994) was purchased from Selleck.

### Cell preparation for BL-Hi-C and library construction

The BL-Hi-C library construction was performed as previously described^19^ with some modifications. In brief, the primary myoblasts were treated with 2% formaldehyde for 15 min (R.T.) to crosslink protein–protein and protein–DNA in the cells, the reaction was quenched by adding glycine solution (F.C. 0.2 M) for 5 min (R.T.). Scrape, transfer and centrifuge to collect the cells, mix cells well and then split the crosslinked cell suspension into aliquots of 3-5 × 10^5^ cells. Cell pellets were frozen in liquid nitrogen and stored at −80°C until use.

The cells were suspended using 1ml 1% SDS lysis buffer (50 mM HEPES-KOH, 150 mM NaCl, 1 mM EDTA, 1% Triton X-100 and 1% SDS) at R.T. for 15 min. Wash the pellet with 1ml 0.1% SDS lysis buffer (50 mM HEPES-KOH, 150 mM NaCl, 1 mM EDTA, 1% Triton X100 and 0.1% SDS). Then centrifuge at 500 g for 2 min, discard the supernatant and add 50 μl of 0.5% SDS for 30 min at 37°C. Add 145 μl H_2_O and 12.5 μl 20% Triton X-100 to quench SDS. Then, the genome was digested by 5 μl enzyme HaeIII (10 U/μl, NEB) with 81.5 μl H_2_O, 10 μl 10 × NEB buffer 2, 2.5 μl 20% Triton X-100 and 1 μl BSA into fragments with blunt-ends at 37°C for at least 2 hrs. Next, the blunt-ends of the DNA fragments were treated with adenine and ligated with bridge linker containing biotin for 4 hrs (at R.T.). Next, add 2 μl 10 mM dATP and 5 μl Klenow Fragment (3’ −> 5’ exo-) (NEB) to digest the unligated DNA fragments at 37°C for 40 min. Then pellet the cells and resuspend with 79 μl H_2_O, 2 μl Bridge Linker, 10 μl T4 DNA ligase buffer, 5 μl 20% Triton X-100, 1 μl BSA and 5 μl T4 DNA ligase to perform the proximity ligation assay at R.T. for 4 hrs. Spin down and add 88 μl H_2_O, 10 μl lambda Exonuclease buffer, 1 μl lambda Exonuclease and 1 μl Exonuclease I to remove unligated Bridge Linker. The cells were then digested by 2 μl 10% SDS, 5 μl 20mg/ml Proteinase K (Ambion) and 4 μl 5M NaCl at 65°C for 6 hrs or overnight. The DNA was purified following phenol-chloroform (Solarbio) extraction and ethanol precipitation. Further, the DNA was fragmented to 300 bp average length using the S220 Focused-ultrasonicator (Covaris), and the biotin-labeled DNA fragments were pulled by the streptavidin coated Dynabeads M280 (Thermo Fisher). Then discard the supernatant and add 78 μl H_2_O, 10 μl T4 DNA ligase buffer, 1 μl 10mM dNTP, 5 μl T4 polynucleotide kinase (NEB), 1 μl T4 DNA Polymerase (NEB) and 5 μl Large (Klenow) fragment (NEB) to the DNA at R.T. for 30 min to repair end. Next, discard the supernatant and add 83 μl H_2_O, 10 μl NEB buffer 2, 2 μl 10 mM dATP and 5 μl Klenow Fragment (3’ −> 5’ exo-) (NEB) for A-tailing at 37°C for 30 min. Then discard the supernatant and add 7.6 μl H_2_O, 10 μl 2 × Quick ligase buffer, 2 μl Quick ligase and 0.4 μl 20 μM Y-adaptor for sequencing adaptor ligation at R.T. for 30min. The library from the beads was then amplified by PCR of 12-14 cycles. After purification with the AMPure XP beads (Beckman, Germany), the libraries were sequenced as double-end 150-bp reads. The sequences of Bridge Linker and Y-adaptor were from the BL-Hi-C paper^19^.

### ChIP and library preparation

Primary myoblasts were fixed with 1% formaldehyde in F-10 at R.T. for 10 min and quenched by adding glycine (F.C. 0.125 M) for 5 min (R.T.). Then the cells were scraped, pelleted, frozen in liquid nitrogen and stored at 80°C until further use. Next, 1 × 10^6^ cells per IP for H3K27ac, 2.5 × 10^6^ cells for CTCF, 1 × 10^7^ cells for MyoD were then thawed on ice, resuspended in cold cell lysis buffer (10 mM Tris pH 8.0, 10 mM NaCl, 0.5% NP-40) + 1 × EDTA-free Protease Inhibitors. Cells were lysed for 15 min on ice with occasional inversion every 2 min and centrifuged to pellet nucleus. Then the nuclear pellet was resuspended with cold nuclei lysis buffer (50 mM Tris pH8.1, 10 mM EDTA, 1% SDS) + 1 × Protease Inhibitors. Nuclei were lysed for 15 min on ice with occasional inversion every 2 min and then sonicated for 24 cycles with 30s on and 30s off using Bioruptor (Diagenode). After sonication, 10 × volumes of IP dilution buffer (20 mM Tris pH 8.0, 2 mM EDTA, 150 mM NaCl, 1% Triton X-100 + protease inhibitors) was added, chromatin was precleared using 25 μl Protein A/G magnetic dynabeads (ThermoFisher, Cat.N: 88802/03) / IP for 1 hr at 4°C with rotation. Then chromatin was incubated with the antibody for 6 hrs or overnight at 4°C with rotation (The following antibodies were used for ChIP: MyoD, sc32758; H3K27ac, ab4729; CTCF, CST2899S). Meanwhile, 25 μl beads / IP were blocked with 1 ml PBS/BSA (1 × PBS + 5 mg/ml BSA-fraction V) at R.T. for 3 hrs or overnight at 4°C. Beads were then added to the chromatin/antibody and incubated 4 hrs at 4°C with rotation. Next, beads were washed twice with cold low salt wash buffer (20 mM Tris pH 8.1, 2 mM EDTA, 150 mM NaCl, 1% Triton X-100, 0.1% SDS), twice with high salt wash buffer (20 mM Tris pH 8.1, 2 mM EDTA, 500 mM NaCl, 1% Triton X-100, 0.1% SDS), twice with cold LiCl wash buffer (10 mM Tris pH 8.1, 1 mM EDTA, 250 mM LiCl, 1% NP-40, 1% sodium deoxycholate) and once with cold 1 × TE buffer (10 mM Tris pH8.0, 1 mM EDTA). DNA: protein complexes were then eluted twice for 10 min at 70°C in 150 μl elution buffer (100 mM NaHCO_3_, 1 mM EDTA, 1% SDS) each time. 5 M NaCl, RNase A and proteinase K were then added and samples + inputs were reverse cross-linked at 65°C for 6 hrs or overnight and DNA was purified using phenol/chloroform extraction and ethanol precipitation. The library construction was performed using NEBNext^®^ Ultra™ II DNA Library Prep Kit for Illumina.

### Total RNA library preparation

Total RNA was extracted using TRIZOL (Ambion, USA). The library construction was performed by ANNOROAD and Novagene. Sequencing libraries were constructed using NEBNext UltraTM RNA Library Prep Kit for Illumina (#E7530L, NEB, USA) following the manufacturer’s recommendations and index codes were added to attribute sequences to each sample.

### Library QC and sequencing

Before sequencing, the libraries were quantified by qPCR and the size distribution was assessed using Agilent 2100 Bioanalyzer. Hi-C libraries were sequenced 2 × 150 bp pair-end on the Illumina Nova seq S2 platform. ChIP-seq libraries were sequenced 1 × 50 bp single-end on the Illumina Nextseq550/Nova seq S2 platforms. RNA-seq Libraries were sequenced 2 × 150 bp paired-end run on the Illumina Nova seq S2/HiSeq X10 platforms. All the sequencing was performed by ANNOROAD and Novagene.

### Recombinant MyoD purification

The PCR-amplified DNA fragments encoding full-length MyoD (residues 1-318) was cloned into a modified pET28a vector with an N-terminal His6-SUMO tag and Ulp1 protease site. The constructed expression vector named pET28a-SUMO-MyoD-FL was transformed into the Escherichia coli strain BL21 (DE3) (Agilent Technologies, Santa Clara, CA, USA). The cells were grown in LB medium supplemented with 50 mg/ml kanamycin at 37°C until OD600 reached 0.6-0.8, and then cultured overnight at 18°C after adding 0.2 mM Isopropyl β-D-1-thiogalactopyranoside (J&K) to induce protein expression.

The cells were harvested by centrifugation at 5000 rpm (Thermo Fisher Scientific) for 15 min. The pellet was resuspended in buffer containing 20 mM Tris–HCl pH 8.0, 500 mM NaCl, 25 mM imidazole pH 8.0 and lysed under high pressure via JN-02C cell crusher and further clarified by centrifugation at 17,000 rpm, 60 min, 4°C (Beckman Coulter). Supernatant was loaded onto 5 ml HisTrap FF columns (GE Healthcare, Beijing, China) pre-equilibrated with Buffer 1 (20 mM Tris-HCl pH 8.0, 500 mM NaCl, 25 mM imidazole pH 8.0). The His6-SUMO-MyoD-full-length (FL) was eluted from the column using elution buffer (Buffer2, 20 mM Tris-HCl pH 8.0, 500 mM NaCl, 500 mM imidazole pH 8.0) with a stage-wise gradient on ÄKTA™pure (GE Healthcare). His6-SUMO tags were cleaved by Ulp1 protease during dialysis against buffer S (20 mM Tris-HCl, pH 8.0, 500 mM NaCl) and removed by a second step HisTrap FF column (GE Healthcare). The MyoD-FL protein in flow through was diluted with pre-cooling 20 mM Tris-HCl pH 8.0 to ensure a low NaCl concentration to remove excess nucleic acids by HiTrap SP FF column (GE Healthcare, Beijing, China) with buffer A containing 20 mM Tris-HCl pH 8.0, 200 mM NaCl and buffer B containing 20 mM Tris-HCl pH 8.0, 1 M NaCl. Eluted protein was concentrated by centrifugal ultrafiltration (Millipore Amicon Ultra, 10K), and loaded onto a pre-equilibrated HiLoad Superdex 75 16/60 column (GE Healthcare) in buffer GF (20 mM Tris-HCl, pH 8.0, 500 mM NaCl, 2 mM DTT) for final purification. All steps should perform on ice or at a low temperature.

### *In vitro* DNA looping assay and Transmission Electron Microscopy (TEM)

First, plasmid pUC57-MyoD was constructed. pUC57-MyoD contains 10 MyoD binding sites (E-box, CAGCTG) separated by ~2.8 kb of intervening DNA. The intervening DNA was chosen as regions without MyoD occupancy or MyoD binding motifs based on ChIP-seq and motif distribution in muscle cells. The MyoD binding motif (CAGCTG) was validated with EMSA (our previous data). Next, the plasmid was digested with EcoRI (NEB R0101) and gel purified.

The ligation assay was carried out as follows. The DNA and proteins were incubated in binding buffer (20 mM Tris 7.9, 50 mM NaCl, 1 mM EDTA) at 25°C for 20 min with a ratio of 1:100, while the final concentration of DNA is 80 nM and the final concentration of proteins is 8 μM. Then, DNA-protein complexes were purified by gravity-flow gel filtration (4 ml of 2% agarose, ABT E-01508S-2B) using TE buffer (10 mM Tris 7.9, 1 mM EDTA). Peak fractions of DNA-protein complexes were analyzed by TEM. Briefly, samples were fixed with 0.4% glutaraldehyde in TE buffer on ice for 30 min. Then, DNA-protein complexes samples were mixed with a buffer containing spermidine to a final concentration of 2 mM to enhance the absorption of the chromatins to the grids. Samples were loaded to the glow-discharged carbon-coated EM grids and incubated for 2 min and then blotted. The grids were washed stepwise in 20 ml baths of 0%, 25%, 50%, 75%, and 100% ethanol solution for 4 min (each at R.T.), air dried and then shadowed with tungsten at an angle of 10° with rotation. Finally, the samples were examined using a FEI Tecnai G2 Spirit 120-kV TEM. Micrographs are shown in reverse contrast. The statistical significance was calculated using the two-tail student’s t-test.

### DNA FISH

DNA FISH probes were constructed according to the bioRxiv preprint (Tn5-FISH)^44^. Briefly, the probe library was generated with PCR amplification and recovery by DNA Cleanup kit (D4014, Zymo research, U.S.A.). Then Tn5-FISH probes were amplified by a second PCR with fluorescence-tagged primers. The *in situ* hybridization procedure of Tn5-FISH was similar with traditional FISH as previously described^45^. Briefly, cells were seeded on coverslips and fixed with 4% paraformaldehyde for 15 min, washed with PBS for 5 min, followed by permeabilization with 0.1% Triton X-100/saponin solution for 10 min. Next, cells were washed with PBS for 5 min and incubated in 20% glycerol for 20 min. The cells were then snap freeze-thawed with liquid nitrogen for three times, air-dried and washed by PBS. Cell were further treated with 0.1 M HCl, permeated with 0.5% Triton X-100/saponin and digested with 100 μg/mL RNase A for 30 min at 37°C, then balanced in 50% deionized formamide/2 × SSC solution for 30 min. Each color of 10 ng Tn5-FISH probe was mixed with hybridization buffer, then added on the slide and coverslip was placed on top, denatured at 75°C for 5 min, then 37°C overnight. The coverslips were washed with wash buffer, stained with DAPI, and then mounted on slides by AntiFade mounting medium. The slides were then imaged by Zeiss LSM780 confocal microscope. The images were processed by FIJI software (version 1.25 h, from NIH) and quantified by Imaris 9.2.0 (Bitplane, Switzerland). The images were then processed with deconvolution software Huygens Pro 17.10. The statistical analyses were achieved with build-in two-tailed t-test models from Graphpad Prism7 software (Graphpad Software, San Diego, U.S.A.).

The primer sequences for probe amplification were as follows:

*Mybph*-1F CCTACTGCTGACCCTTAAACACTCC,
*Mybph*-1R CCAGCCTCCATCCATCACCTCAAG,
*Mybph*-2F CTAGTCAAGGCCTCTAGGAGGAG,
*Mybph*-2R GATGCACGGTGAGCTCATAGCG,
*Mybph*-3F GAAGCCATACTCTTTAACATCAGCCAG,
*Mybph*-3R GGGTAGCACATTGTGAATCAGAAAGC,
*Mybph*-4F GACACACCATCTCTGGGACTTTCC,
*Mybph*-4R AGTTCGAGAAGGTTATCAGCATGGAG;
*Myog*-1F GCTACTGCCCTCAACTGTAGC,
*Myog*-1R CTGTGTCCTGTGCATTGGGTAC,
*Myog*-2F TCTGCCACCTCATTGGCTACAAC,
*Myog*-2R GCTCTTAATCATCTCTTCTGTGCT,
*Myog*-3F ATCTGGAGTGGTCCTGATGTGG,
*Myog*-3R GTGGCTATCCCTAGGTGAAGCC,
*Myog*-4F CACTCCTGAGTCTTCCTTACCCA,
*Myog*-4R TGAATCCCTGTCTGCAATGGTTTG.

### Immunofluorescent staining

Cells were fixed in 4% paraformaldehyde in PBS, blocked in 3% BSA and stained according to standard protocols using primary antibodies against MyoD (santa cruz, 1:500), CTCF (millipore, 1:200), Pax7 (DSHB, 1:15), MyoG (DSHB, F5D, 1:500), MHC (DSHB, MF20, 1:500) and Alexa Fluor dye conjugated secondary antibodies (zhongshanjinqiao) and DAPI. Secondary antibodies used for dSTORM: Goat anti-mouse Alexa Fluor 488 (Life Technologies, A-11017) and goat anti-rabbit Alexa Fluor 647(Life Technologies A-21246).

### Direct stochastic optical reconstruction microscopy (dSTORM)

Coverslips (Warner Instruments) with 25 mm 1.5 thickness were used. C2C12 cells were cultured on cleaned coverslips coated with fibronectin (2 μg/mL in PBS, pH=7.4; Sigma). Following immunostaining of MyoD and CTCF (See immunoflueroscent staining section above), cells were blocked with 1 × PBS solution containing 50 mM Glycine for 10 min twice and then washed once in 1 × PBS before being subjected to dual-color imaging in dSTORM imaging buffer (1 × PBS, 100 mM β-mercaptoethylamine (Sigma), pH 7.4) containing Tetraspek beads (Life Technologies), which were used as fiducial markers for x-y drift correction and for overlaying two color images.

dSTORM images were acquired on an Olympus IX83 motorized inverted fluorescence microscope equipped with the CellTIRF-4Line system, a 150 × 1.45 NA total internal reflection objective, a back-illuminated EMCCD camera (Andor), the IX3-U-m4TIRSbx filter set (Olympus) and 488 nm (150 mW) and 640 nm (140 mW) lasers. dSTORM images of Alexa 488 and Alexa 647 were acquired sequentially, with the acquisition of Alexa 647 preceding the acquisition of Alexa 488. All images were acquired at 10 frames per second. Individual molecules were localized and super-resolution images were reconstructed using the ThunderSTORM analysis module available for ImageJ.

### RNA-seq data processing and differential gene expression analysis

Paired-end RNA-seq reads were obtained from three biological duplicates of WT and MKO primary myoblasts at proliferation (GM) and early differentiation (DM) stage, with an average read depth as 4.5 × 10^7^ read pairs per sample. HISAT2 v2.1.0^46^ was used to align the reads to mm9 genome, then HTSeq v0.6.0^47^ was applied to calculate the counts per gene with default parameters. Differential genes were identified with FDR <0.01 and foldchange>1.5 by DESeq2 v1.24.0^48^.

### ChIP-seq data analysis

H3K27ac and CTCF ChIP-seq analyses were performed with two replicates in WT and MKO primary myoblasts at proliferation (GM) and early differentiation (DM) stage while MyoD ChIP-seq analyses were in WT myoblasts and myocytes. H3K27ac ChIP-seq analyses were also performed in MKO cells with or without Tacedinaline treatment. ChIP-seq reads were mapped to the mm9 genome with Bowtie2 v2.3.5^49^ using default parameters. Aligned reads were filtered with minimum MAPQ of 20, and duplicates were removed using SAMtools v1.9^50^. Peaks were called using MACS2 v2.2.5^51^ with default parameters for CTCF and MyoD.

H3K27ac peaks were called by MACS2 with broad peak mode. Signal tracks were generated by using the-SPMR option in MACS2. Then, the UCSC Genome Browser utility^52^ bedGraphToBigWig was used to transform the bedgraph files to bigwig files. Differential Peaks of ChIP-seq experiments were called with R package DiffBind v2.10.0^53^ with default settings. Heatmaps of ChIP-seq signal enrichment were generated by python package deeptools v3.0.1^54^. ChIP-seq peak annotation were done by ChIPseeker v1.20.0^55^. Public ChIP-seq data used in this study were processed in the same manner. Specially, we merged our detected MyoD peaks with those called from public MyoD ChIP-seq data (GSE56131). Then, the differential peaks were determined by examining the MyoD binding strength on merged peaks comparing our WT-DM and WT-GM Myod ChIP-seq data based on DiffBind v2.10.0. WT-GM pseudo-peaks combined static MyoD peaks and MyoD peaks with reduced signal in WT-DM while WT-DM pseudo-peaks combined static MyoD peaks and MyoD peaks with enhanced signal in WT-DM cells.

### Motif enrichment analysis

The homer^56^ script findMotifsGenome was used with default parameters for enriching the motif of ATAC-seq peaks overlapped with loop anchors in WT primary myoblasts. The homer script findPeaks with option-nfr was used to find the nucleosome free regions in H3K27ac peaks. Then, these nucleosome free regions in broad H3K27ac peaks were used to call motif by findMotifsGenome with size as 200bp.

### Hi-C data processing, Hi-C loop calling and differential loop detection

BL-Hi-C data were processed for WT and MKO primary myoblasts at proliferation (GM) and early differentiation (DM) stage as well as MKO cells with or without Tacedinaline treatment. Non-redundant uniquely mapped contacts were generated utilizing the in house HiCpipe pipeline, which trims the bridge linker, aligns reads, filters artifact fragments, and removes duplicates. Juicer^26^ was used for following Hi-C map generation, compartment partition and loop calling. For every 100 kb bin, A or B compartments were defined by the over 70% replicates majority rule. Contact domain boundaries (CDBs) and their relative insulation scores were calculated by HiCDB while insulation scores^57^ were calculated on MyoD or CTCF bound CDBs with insulation score method (parameters as -is 1000000 -ids 200000 -im mean -nt 0.1). Differential CDBs were defined with a t-test FDR<0.01 and a difference between cases and controls higher than 80% quantile of the overall difference. The domain score^38^ (D-score) for each loop was calculated by dividing the intra-loop interactions with all interactions connected to the corresponding loop. Loops with differential internal interactions (D-score) were determined with a t-test FDR<0.01 and fold change higher than 80% quantile of the overall fold changes.

Loops were called under 3 kb, 5 kb and 10 kb resolution with HiCCUPS using parameters (-p 6,4,2 -i 10,7,5) and scaled to 5 kb and 10 kb resolution. These parameters were also used to determine the loop strength of requested loops with HiCCUPS. Loop strength was calculated as relative Hi-C contacts at loop pixels compared with surrounding Donut-shaped background pixels. The differential loop detection method was adapted from Douglas et al.^40^ which depended on DESeq2 to detect differential enrichment of the loop over background across condition. For comparing loops in two conditions, loops were called on combined BL-Hi-C matrixes for each condition and merged together at first. Then, the loop pixels and their surrounding pixels were collected according to the previous work from each biological replicate at both 5 kb and 10 kb resolution. As previous study described bias^39^ based on contact distance associated with the identification of differential interactions from Hi-C-like data, loops were split into two distance regimes of greater or less than 150 kb for 5 kb and 10 kb resolution. Differential loops called from each set were thresholded by FDR as 0.1 and were combined to form the final set of dynamic loops. For dynamic loops detected under both 5 kb and 10 kb resolution, 5 kb resolution loops were kept. Public *in situ* Hi-C data of *in vivo* ES and CN were processed by HiC-Pro v2.11.1^58^ in the mapping and filtering step and then treated with the same processing framework as our BL-Hi-C data.

### Loop annotation, aggregation and functional analysis

Loop anchors (scaled to 10 kb) were overlapped with MyoD and CTCF binding sites and classified into four classes using bedtools toolset v2.27.1^59^. MyoD or CTCF occupancy on loops were also determined by intersecting loop anchors and MyoD or CTCF peaks. Aggregate peak analysis (APA) plots were generated to assess the quality of loop detection and explore the characteristics of different loop classes by Juicer APA command. The aggregated maps generated by Juicer were normalized by the actual supporting loop number and the cis interaction pairs of each sample. The APA score was also calculated by Juicer APA command as the ratio of Hi-C contacts at the central pixel to the mean Hi-C contacts of the lower left pixels. Enrichment plots for differential CDBs and loops with differential internal interactions were generated with coolpup.py^60^. To gain knowledge about the function of different loop types, the loop anchors were analyzed by GREAT v3.0.0^61^ with the association rule: single nearest gene and 100kb max extension to identify the enriched biological process. Specially, for loop internal interaction analysis, we examined the genes enclosed in the loops and the genes were assigned to their most dynamic loops to get the explicit gene-loop pair. In other sections, genes associated to loops were determined if their promoters (±3 kb around TSS) overlapped loop anchors. To measure the muscle lineage specificity of genes associated with MyoD-bound and CTCF-bound loops, we calculated the Z-score of gene expression in WT-DM cells compared with other ENCODE collected cell types (embryonic stem cell, spleen, B cell, T cell, megakaryocyte, neural progenitor and cortical neuron). ClusterProfiler v3.12.0^62^ was used to enrich and compare GO terms of different sets of loops bound by CTCF or MyoD.

### Visualization

Tracks of Hi-C maps and ChIP-seq data were generated by pyGenomeTracks v3.1.2^63^. Hi-C maps of each condition were normalized by cis interaction pairs.

### Statistics

Analysis-specific statistics were applied as described in each subsection. For comparisons of distributions, Wilcoxon rank sum test was employed in R. The statistical significance of the other data was calculated using the two-tail student’s t-test. The following annotation applies for all figures: *P < 0.05, **P < 0.01, ***P < 0.001.

### Code availability

Custom scripts described in the Online Methods will be made available upon request.

### Data availability

The raw sequence data reported in this paper have been deposited in the Genome Sequence Archive (GSA)^64^ under accession number CRA002490 that are publicly accessible at https://bigd.big.ac.cn/gsa. The processed data of this paper have been deposited in the Gene Expression Omnibus (GEO) database under the accession number: GSE157339. Accession codes of the published data in GEO used in this study are as follows: MyoD ChIP-seq of wild-type primary myoblasts, GSE56131; ATAC-seq of wild-type primary myoblasts, GSE63573; NeuroD2 ChIP-seq of embryonic cortical neuron cells, GSE67539; Hi-C data of mouse neural development as well as CTCF and H3K27ac ChIP-seq of embryonic cortical neuron cells, GSE96107; SMC3 ChIP-seq of C2C12 myoblast and myotube, GSE113248.

---

**Extended Data Fig. 1.**
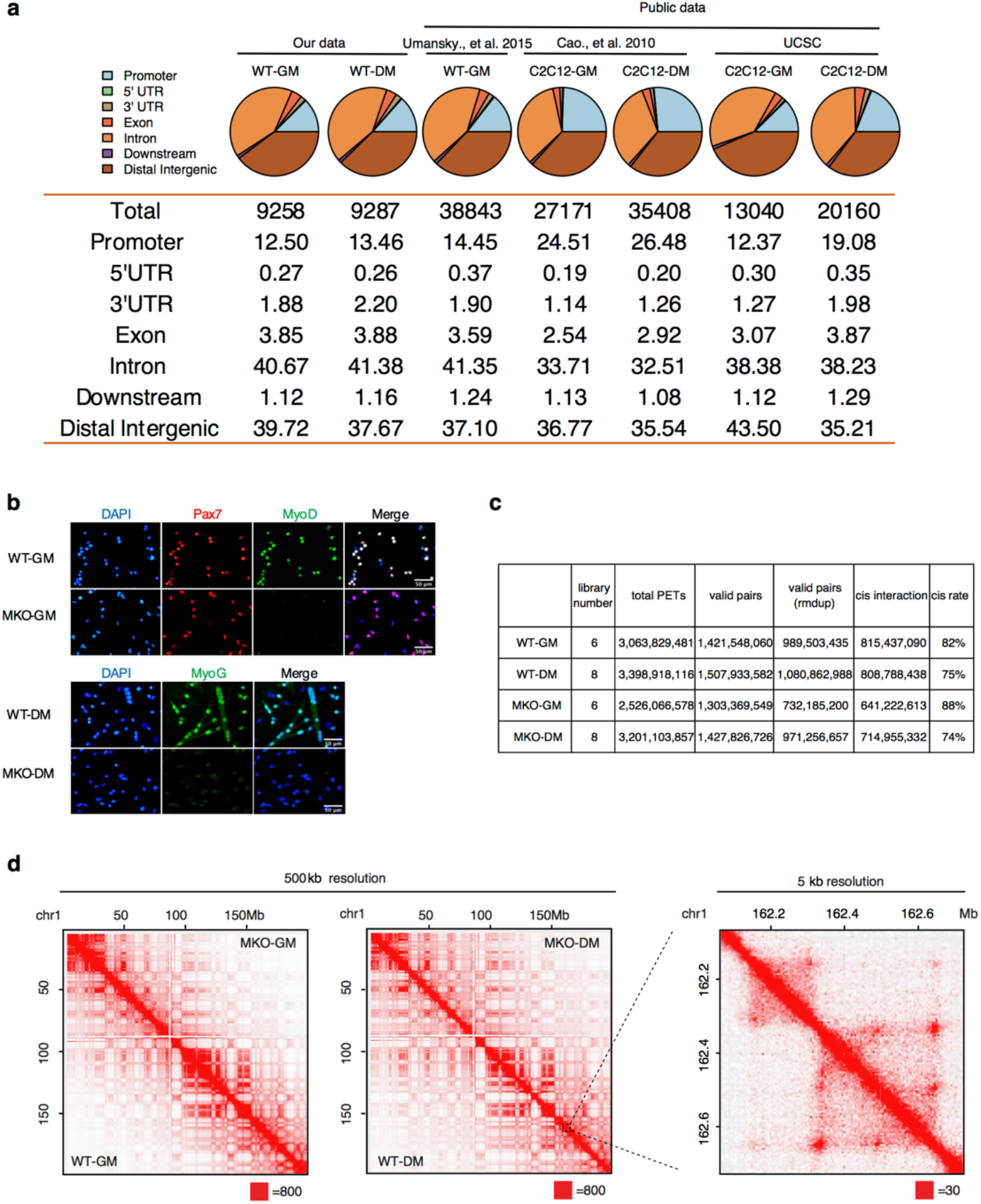
Unveiled genome architectural functions of MyoD in muscle cells. **a**, Genomic distribution of MyoD binding peaks in proliferating or differentiating muscle cells. WT represents primary muscle cells while C2C12 is a skeletal muscle cell line. MyoD ChIP-seq data were from our study and public datasets of indicated references. Promoter is defined as ± 3 kb from the transcription start site (TSS). **b**, Representative images for immunofluorescent staining of Pax7 (red), MyoD (green) and MyoG (green) in wild type (WT) and MyoD knockout (MKO) proliferating muscle stem cells (myoblasts, cultured in growth media, GM) was shown at upper panel. Corresponding data for differentiating muscle stem cells (myocytes, induced to differentiation in differentiation media, DM) was shown at lower panel. DAPI (blue) served to visualize nucleus. Scale bars represent 50 μm. **c**, Depth and quality of sequencing data for BL-Hi-C libraries on four types of samples (WT-GM, WT-DM, MKO-GM and MKO-DM). **d**, Hi-C map showing BL-Hi-C data in our study reached 5 kb resolution. Left heatmaps showing chromatin contact matrices from the whole chromosome 1 (chr1) at 500 kb resolution in each of four cell samples. Right panel was a zoom-in heatmap for chr1: 162 ~ 162.8 Mb at 5 kb resolution. Maximum intensity was indicated in the lower right of each panel.

**Extended Data Fig. 2.**
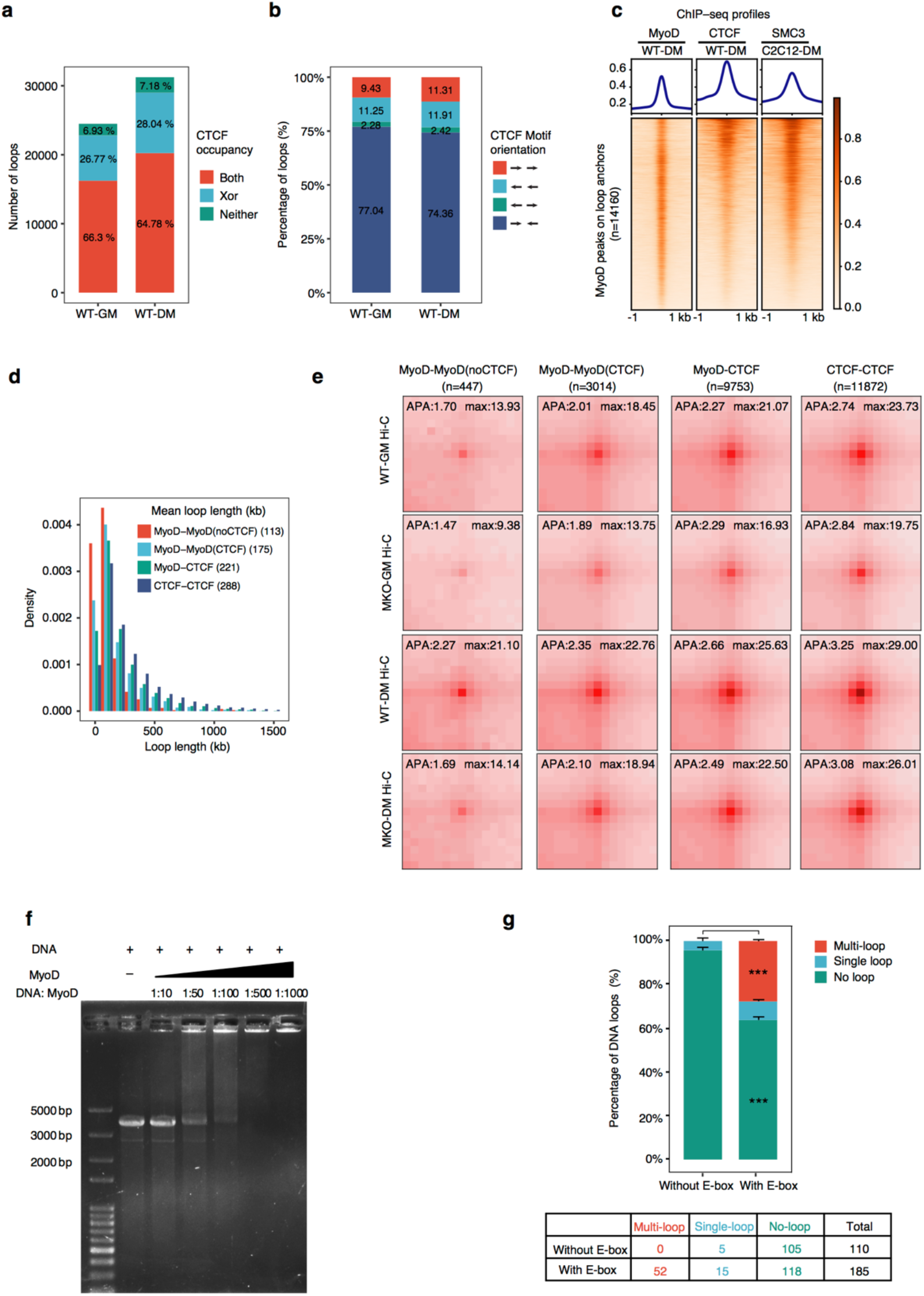
MyoD mediates chromatin loop formation *in vivo* and *in vitro*. **a**, Percentages of chromatin loops with CTCF bound at both anchors (Both, red), one of two anchors (Xor, blue) or neither of two anchors (Neither, green) in WT-GM and WT-DM cells. **b**, Distribution CTCF motif orientation on loop anchors bound by CTCF in WT-GM and WT-DM cells. **c**, Heatmaps of ChIP-seq signal showing the enrichment of MyoD, CTCF and SMC3 on MyoD-bound loop anchors, indicating the concordant binding of MyoD with CTCF and SMC3 on these loop anchors. MyoD and CTCF ChIP-seq were conducted in our study. ChIP-seq for SMC3 in C2C12 cells were referenced from public database (GSE113248). **d**, Loop length distribution of MyoD-MyoD(noCTCF), MyoD-MyoD(CTCF), MyoD-CTCF and CTCF-CTCF chromatin loops. The average loop length was shown at upper right corner. **e**, APA plot showed the aggregated Hi-C contacts around MyoD-bound three types of loops and CTCF-CTCF chromatin loops in MKO cells compared to WT cells. All four type of samples from GM and DM stages were shown. n represents number of each type of chromatin loops. **f**, Free and the recombinant MyoD protein-bound DNA fragment (3.7kb, with E-box) were shown and separated by polyacrylamide gel electrophoresis (PAGE). The binding of MyoD and DNA fragment also showed ratio-dependency. **g**, Statistics of the percentage and number of loops observed with DNA circularization in Fig. 2h. ***P < 0.001, two-tail student’s t-test.

**Extended Data Fig. 3.**
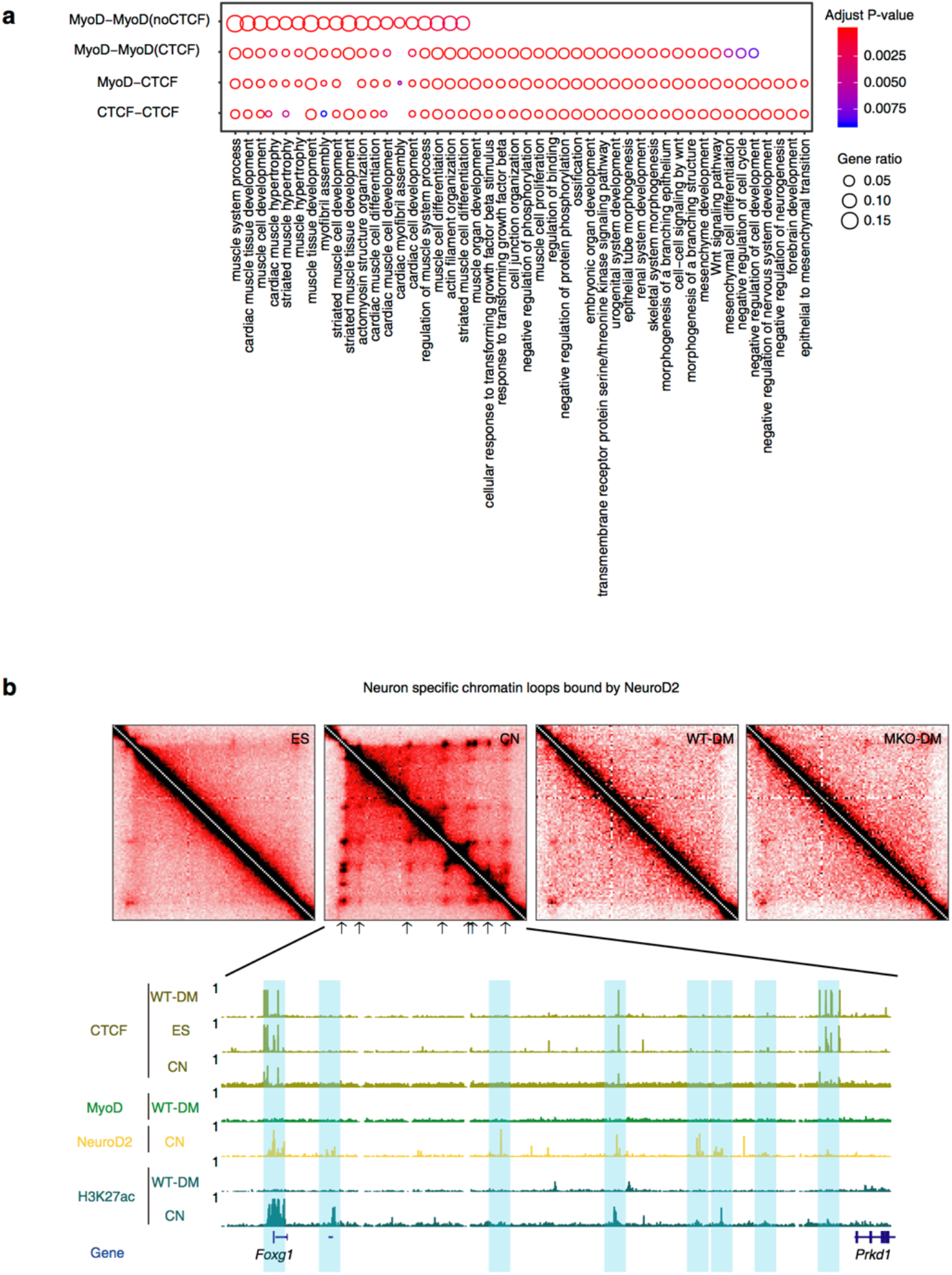
MyoD and NeuroD2 instructs lineage specific chromatin loops. **a**, Enriched GO terms of genes with promoter overlapped with anchors of MyoD-MyoD(noCTCF), MyoD-MyoD(CTCF), MyoD-CTCF and CTCF-CTCF chromatin loops. **b**, Representative region of neurogenic lineage specific chromatin looping at *Foxg1* locus bound by NeuroD2.

**Extended Data Fig. 4.**
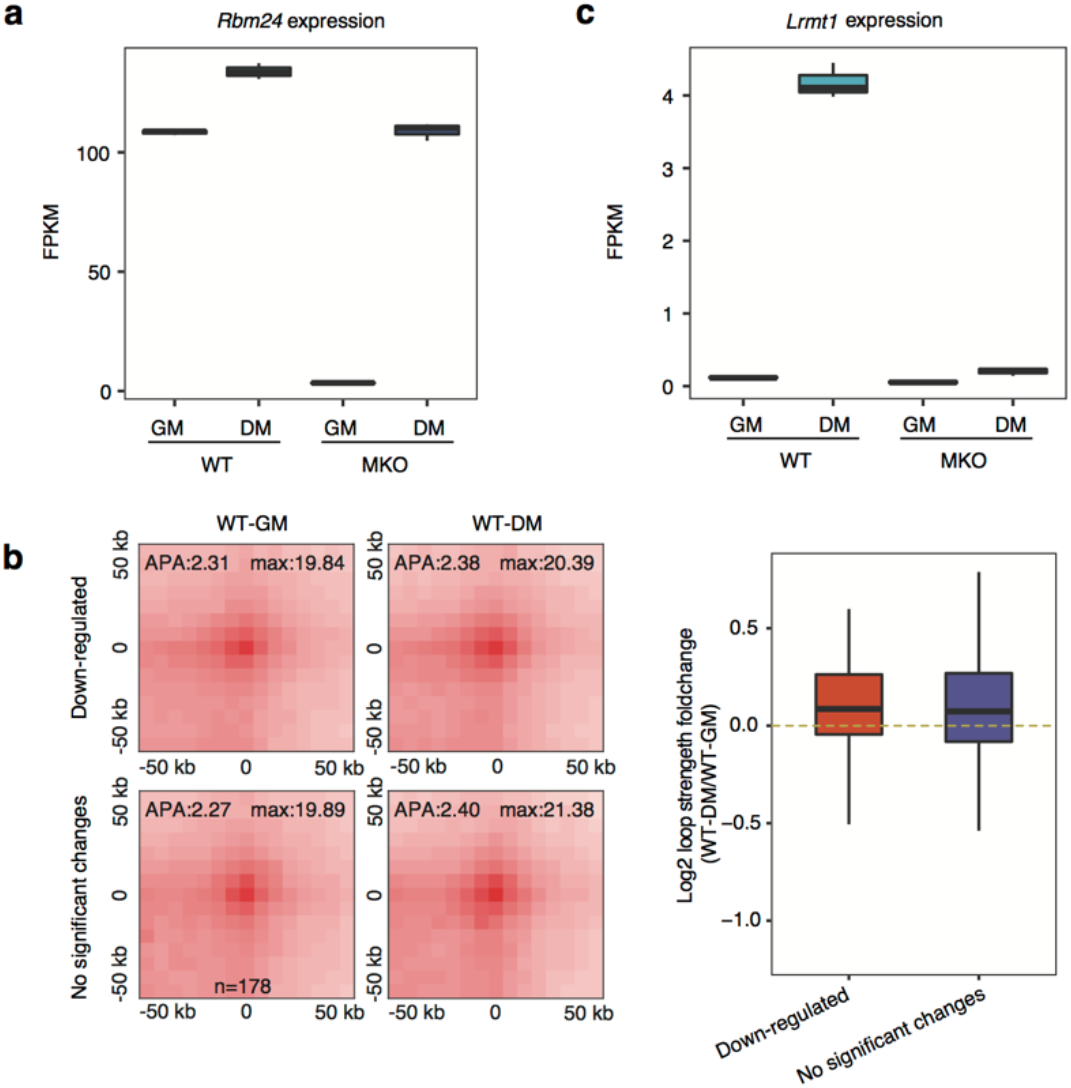
MyoD mediates primed architectural loops in myoblasts. **a**, Boxplot showing the gene expression (FPKM) of *Rbm24* in four types of samples (WT-GM, WT-DM, MKO-GM and MKO-DM). **b,** Loop changes of MyoD-bound architectural loops during differentiation. APA plot showed the aggregated Hi-C contacts around internal interaction reduced MyoD-bound chromatin loops with down-regulated genes or not significantly changed genes in WT-GM and WT-DM cells. Boxplot showed the loop strength fold changes for loops with down-regulated genes or not significantly changed genes between WT-DM and WT-GM cells. **c,** Boxplot showing the gene expression (FPKM) of *Lrmt1* among four types of samples (WT-GM, WT-DM, MKO-GM and MKO-DM).

**Extended Data Fig. 5.**
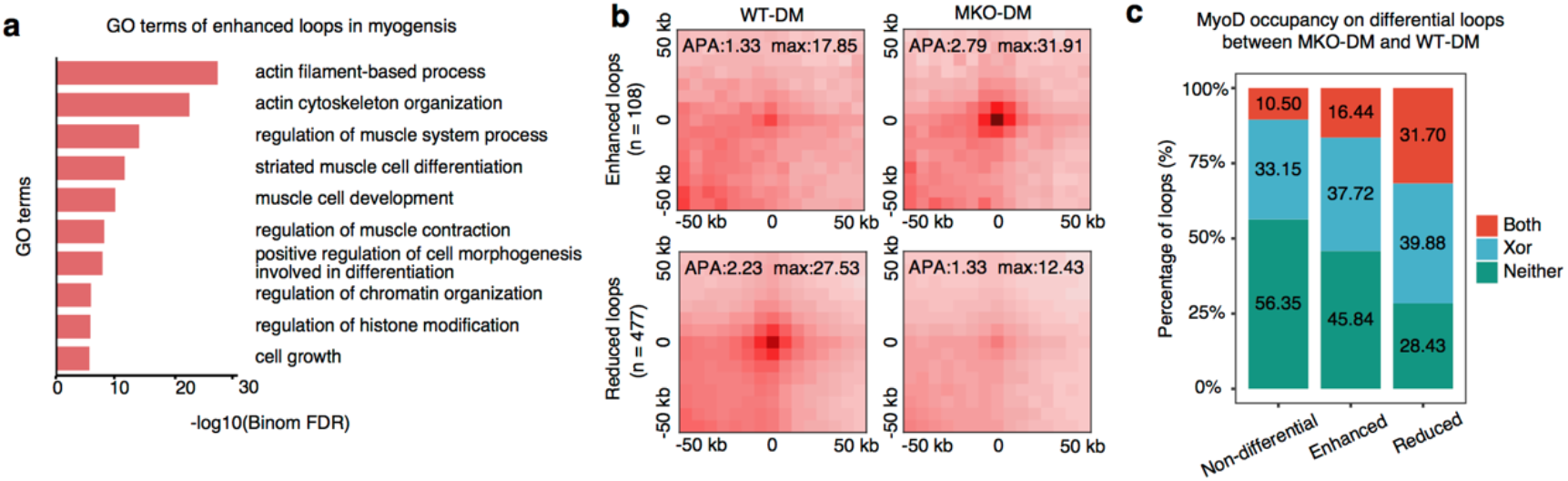
MyoD-induced regulatory loops are functionally required for muscle cell differentiation. **a**, Enriched GO terms of enhanced loops in WT-DM cells compared with WT-GM cells. **b,** APA plots showing aggregated Hi-C contacts around significantly reduced or enhanced chromatin loops in MKO-DM cells compared to WT-DM cells. n represents number of the differential chromatin loops between MKO-DM and WT-DM cells. Differential loops were identified with DESeq2 (Methods). **c,** Percentages of MyoD-bound loops in the non-differential, enhanced or reduced chromatin loops described in panel b. MyoD bound at both anchors (Both, red), one of two anchors (Xor, blue) or neither of two anchors (Neither, green) were shown. Pseudo-peaks of MyoD binding with combined ChIP-seq data from public and our study were used for this analysis.

**Extended Data Fig. 6.**
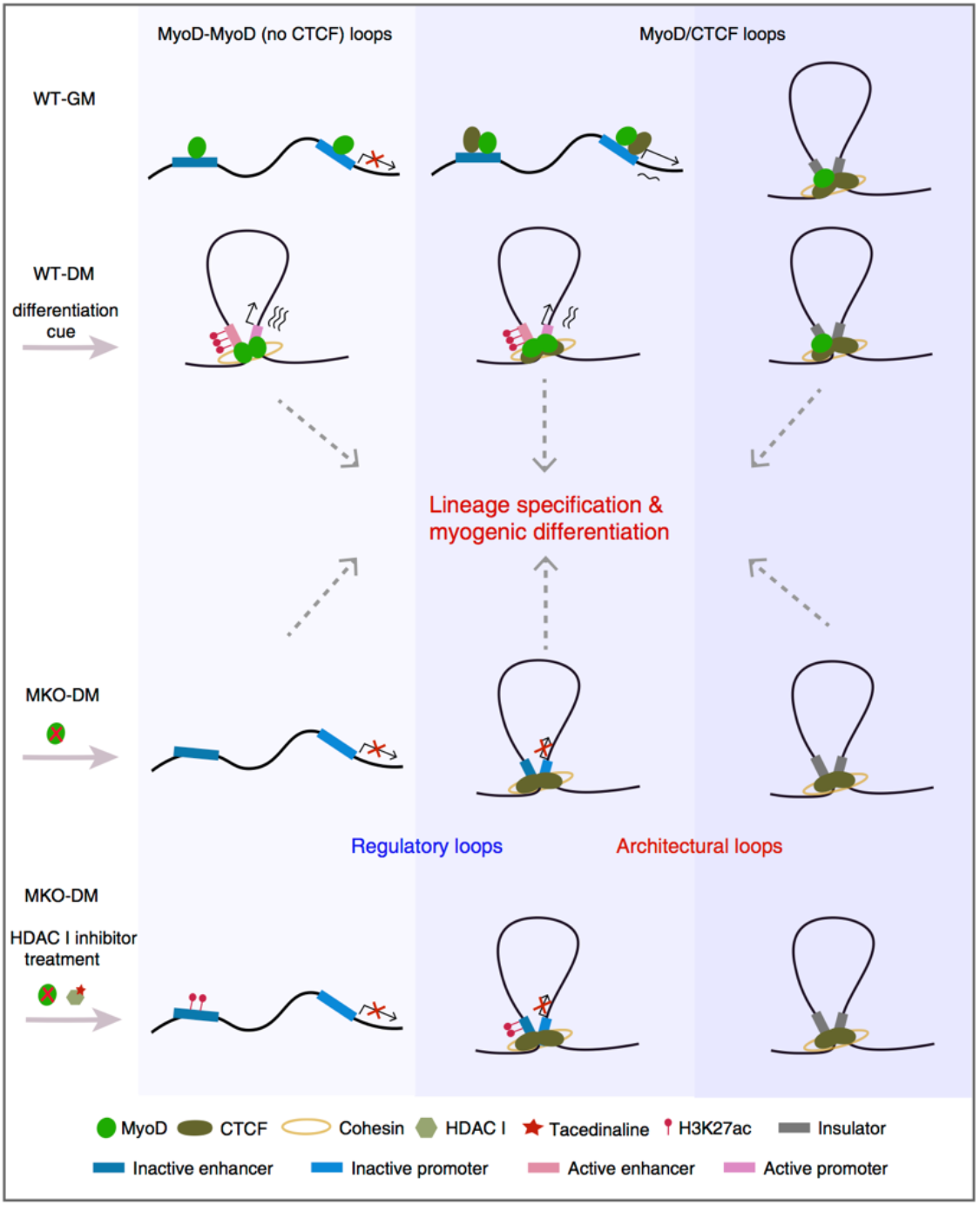
Hypothetical model for MyoD functions as a structure organizer of 3D genome architecture in muscle cells. MyoD is expressed in proliferating cells and functions as genome organizer that specifies the proper 3D genome architecture unique to muscle cell via two distinct mechanisms: acting directly by itself and through interactions with CTCF protein. Intriguingly, the MyoD-bound loops including pre-existed in proliferating cells and differentiation-induced play dual roles of regulatory and architectural in differentiating cells, which underscore a novel idea that a given cell type specific pioneer TF may in fact hide architectural potential.

